# Modular Cell-Free Extract Mixing Enables Soluble, Active Biomanufacturing of Difficult-to-Express and Aggregation-Prone Proteins

**DOI:** 10.64898/2026.10.09.757855

**Authors:** Larissa Firmansyah, Anabella Delmundo, Javin P. Oza

## Abstract

Cell-free protein synthesis (CFPS) is a powerful platform for rapid protein prototyping and biomanufacturing, but expression of complex proteins remains limited by labor-intensive extract preparation as well as the inability of conventional *Escherichia coli* extracts to efficiently support disulfide bond formation and prevent protein aggregation. We previously developed a Cell-Free Autoinduction (CFAI) workflow that simplifies extract preparation, however, CFAI-derived extracts retain the folding limitations of standard *E. coli* systems for producing traditionally difficult proteins. Here, we address these challenges by utilizing extracts enriched with molecular chaperones or Erv1p-based oxidative folding catalysts and combining them through a modular extract mixing strategy to enable independent tuning of the cell-free folding environment. This workflow supplies folding factors through enriched extracts, avoiding recurring purchases of purified supplements and enabling a modeled reagent-cost advantage. To evaluate real-world applications and utility of lysate mixing for protein production, we demonstrate a 10-fold increase in solubility of the aggregation-prone bacterial toxin Colicin M, achieving >95% soluble protein while retaining antimicrobial activity. We further used a Design of Experiments (DoE) framework to optimize combinations of chaperone-enriched and Erv1p-based oxidative folding extracts containing either protein disulfide isomerase (PDI) or DsbC. The optimized compositions produced >95% soluble truncated human tissue plasminogen activator (vtPA), comprising the kringle 2 and protease domains, while preserving enzymatic activity. Together, these results demonstrate that CFAI-derived modular extract mixing provides a tunable, high-throughput strategy with low modeled materials costs for tailoring cell-free biochemical environments for difficult-to-express proteins, expanding CFPS utility for protein engineering and biomanufacturing.

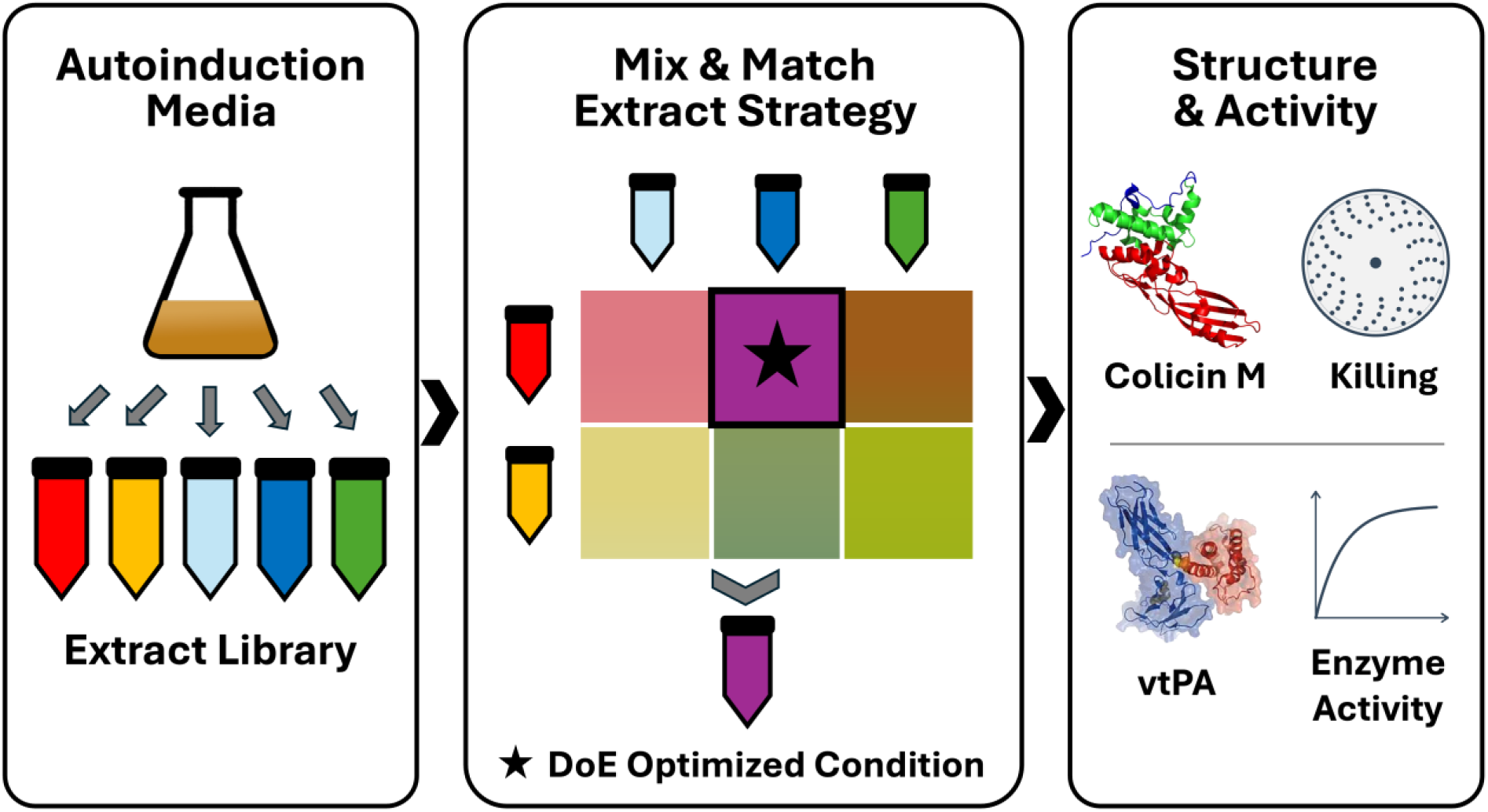

## 1.0 Introduction

Cell-free protein synthesis (CFPS) harnesses transcription and translation machinery along with energy metabolism *in vitro*, moving protein production out of the living cell and into an open reaction that can be directly monitored and modified. Because the reaction is not constrained by host viability, its environment can be manipulated component by component, and all system energy can be focused on production of the protein of interest. The tunable reaction environment enables production of complex or “difficult” proteins whose recombinant expression in *E. coli* may be limited by insoluble or inactive products [1, 2].

The increasing awareness and acceptance of CFPS as a discovery and biomanufacturing platform has been accelerated by improved access to cell-free systems by broader commercial offerings as well as more accessible protocols for preparing reagents in-house. While extract based CFPS systems have been commercially available for decades, an expanding range of commercial offerings has increased access to the platform, with reagent costs dependent on formulation, package size, and folding-factor requirements. Researchers will continue to require tunable systems optimized for new advancements of the platform and bespoke applications, necessitating in-house production of extracts. In order to make in-house preparations more accessible, we previously developed the Cell-Free Autoinduction (CFAI) method, which enables consistent, high-quality extract preparation with significantly reduced technical burden [3]. The key innovation was enabling robust extracts from high-optical density cultures. Combined with autoinduction, this eliminated user intervention between inoculation and harvest. Beyond reducing labor and human error, harvesting at high density also produced a >3× increase in cell mass and therefore extract volume, tripling the extract output for a given batch [4, 5]. The reformulation leading to CFAI also provided new insights, demonstrating that ribosomes were not the limiting factor in producing high-quality extracts from high-OD cultures. Rather, metabolomic analysis suggested that CFAI-based extracts overcame limitations in energy metabolism and redox balance, enhancing overall extract performance from high-OD cultures [3].

Despite these advantages, CFAI performance has only been benchmarked with superfolder GFP (sfGFP), a robust, monomeric reporter that folds readily in the reducing *E. coli* cytoplasm and places minimal demands on the folding machinery. Reporters of this kind are poor proxies for therapeutically and industrially relevant targets that motivate cell-free biomanufacturing including antibodies, proteases, and toxins whose recombinant expression is limited by aggregation or by a requirement for native disulfide bonds. Whether CFAI-derived extracts can express such difficult proteins, and how their folding environment might be tuned to do so, remains uncharacterized. Soluble, active expression of these proteins requires two coupled processes: redox chemistry that forms and rearranges disulfide bonds, and biological chaperone capacity that keeps folding intermediates in solution. Existing approaches address these needs through chaperone supplementation, chaperone-enriched chassis strains, or engineered oxidative folding strains (Sections 1.1 and 1.2), but each fixes at least part of the folding environment within the source strain, whereas an open cell-free reaction allows chemical and biological components to be combined and dosed independently. Here we report an expansion of the CFAI platform using a modular extract mixing strategy augmented by Erv1p-based oxidative folding to enable the soluble expression of therapeutically relevant proteins that remain difficult to produce in traditional *E. coli*-based expression systems. We demonstrate this approach with Colicin M, an aggregation-prone bacterial toxin, and a truncated variant of human tissue plasminogen activator (vtPA), a therapeutic protein that requires nine disulfide bonds, and show that both are produced in soluble form with retained activity. Because the approach builds on the accessible CFAI workflow and combines extracts by simple mixing, it could provide a low-barrier route for matching the folding environment of in-house cell-free reactions to a target protein, extending the reach of cell-free biomanufacturing to difficult toxins and disulfide-bonded therapeutics.

### 1.1 Overcoming Protein Folding Limitations Through Modular Extract Mixing

Two distinct failure modes limit the expression of difficult proteins in *E. coli* extracts. The first is aggregation, in which recombinant proteins misfold when their native folding requirements are mismatched with the chaperone and folding machinery native to the chassis expression system, leaving them aggregation-prone. Chaperone systems such as GrpE/ClpB, DnaK/J/GroESL, and IbpA/B stabilize non-native intermediates and prevent off-pathway aggregation [2, 6].

Chaperone-enriched extracts have improved colicin M solubility in CFPS [7], and purified chaperones can be added directly to cell-free reactions to tune their identity and stoichiometry [8]. We compare these strategies in Section 2.8. Alternatively, chaperones can be overexpressed in the chassis strain during growth, resulting in extracts enriched in the target chaperone without requiring purified chaperone supplementation [7, 9]. Unlike supplementation, chaperone identity and stoichiometry in chassis-enriched extracts are fixed by the growth conditions, becoming untunable parameters, diminishing the advantages of a modular platform technology.

In order to leverage the respective advantages of modular chaperone supplementation and extract enrichment methods while overcoming their limitations, we evaluate whether the chaperone-enriched extract itself can be utilized as the modular, interchangeable component that can be supplemented. The mixing of extracts to augment and complete biochemical machinery is motivated by success in cell-free metabolic engineering, but has surprisingly remained underutilized for protein expression applications [10, 11]. The approach was first demonstrated by mixing enzyme-enriched extracts to reconstruct biosynthetic pathways without re-engineering living organisms and has since been extended to glycoengineering and genetic code expansion [11, 12]. Mixing chaperone-enriched extracts derived from *E. coli* BL21*DE3 offers a route to combine expression efficiency with folding capacity while remaining modular and interchangeable, enabling combinatorial screening without recurring purchases of purified folding factors; the modeled materials costs are evaluated in Section 2.8. Here, we apply modular extract mixing systematically toward the soluble expression of aggregation-prone and difficult-to-express proteins [7, 9].

Toward filling this gap, we utilize *E. coli* extracts enriched with specific chaperone systems: GrpE/ClpB, DnaK/J/GroESL, or IbpA/B [6, 7, 9]. We benchmark the modular extract mixing strategy with Colicin M, a bacterial toxin whose expression in standard cell-free systems is limited by aggregation and poor solubility. Colicin M was selected as a difficult-to-express protein because it lacks disulfide bonds, allowing us to isolate the contribution of chaperone supplementation from oxidative folding [2, 7].

### 1.2 Overcoming Redox Limitations in the E. coli Cytoplasm

The second failure mode for protein expression in cell-free systems, disulfide bond formation, lies beyond the reach of chaperone supplementation alone and remains a central obstacle for most E. coli-based extracts [13, 14]. Although *E. coli* does have the ability to form disulfide bonds, its native oxidative folding catalysts (DsbC) are expressed in the periplasm, leaving the cytoplasm and the resulting cell extract devoid of the proper machinery for oxidative protein folding [15, 16]. The reducing cytoplasmic environmentdisfavors disulfide bond formation, leading to poor stability and incorrect folding, ultimately leading to protein aggregation. This limitation is especially consequential because many proteins of interest in biotechnology and medicine are eukaryotic and contain multiple disulfide bonds that are essential for proper structure and function [15, 16, 17, 18].

Solutions available for cell-free expression can be informed by *in vivo* expression systems. Two strategies have emerged to enable disulfide bonded protein production in *E. coli* for in vivo expression. One widely used strategy genetically disrupts reducing pathways such as thioredoxin reductase (trxB) and glutathione reductase (gor), as in SHuffle strains [19, 20].These engineered strains support cytoplasmic expression of disulfide-bonded proteins [20]. Antibody expression has also been demonstrated in a cell-free platform [21].This strategy has been adapted to cell-free systems. Extracts prepared from the SHuffle T7 strain have been used to support disulfide-bonded protein synthesis in vitro [22].Alternatively, the open nature of cell-free systems uniquely enables an alternate approach in which standard extracts can be supplemented with purified oxidative folding catalysts, such as DsbC or PDI to promote disulfide bond formation without an engineered source strain [13, 14, 21, 23]. Extracts from specialized strains as well as supplementation have proven to be successful paths, each with key strengths and challenges.

The second strategy for *in vivo* expression augments *E. coli*’s native machinery with oxidative folding catalysts into the cytoplasm, as in the Cytoplasmic Disulfide bond formation in *E. coli* (CyDisCo) system [24, 25]. The addition of oxidative folding catalysts, specifically the sulfhydryl oxidase Erv1p co-expressed with either the disulfide isomerase PDI or DsbC (hereafter Erv1p–PDI and Erv1p–DsbC, or collectively the Erv1p–isomerase system), improves functional protein expression by coupling de novo disulfide bond formation with rearrangement of incorrectly paired disulfides during folding [24, 25]. In a direct comparison across fourteen disulfide-bonded proteins, CyDisCo produced higher purified yields than SHuffle for most targets and yielded higher-quality protein, although the performance difference was reduced in rich autoinduction media and was most apparent for proteins with strong oxidative-folding requirements [15]. Notably,the Erv1p-based oxidative folding system enables disulfide bond formation without disrupting reducing pathways, thereby maintaining native redox balance and reducing growth defects [24, 25]. Erv1p acts as the primary driver of disulfide bond formation, transferring electrons from cysteines to molecular oxygen [24, 25]. To facilitate correct disulfide bond formation, disulfide isomerases (PDI or DsbC) rearrange non-native disulfides.

Cytoplasmic expression of human PDI mimics the folding environment of the eukaryotic endoplasmic reticulum (ER), making it particularly effective for complex human proteins. Alternatively, the expression of prokaryotic DsbC replicates the oxidative proofreading that occurs in the *E. coli* periplasm, where secreted proteins are shuttled and isomerized [15, 26]. The Erv1p–isomerase system expresses oxidative folding components directly in the cytoplasm to enable *E. coli* to produce complex, disulfide bonded proteins while maintaining native cellular physiology [24, 25].

The two strategies, Shuffle vs CyDisCO, present complementary tradeoffs. Genetic disruption yields a stable, heritable oxidizing cytoplasm but constrains expression to a specialized strain and imposes a fitness cost on the host [15, 20]. Augmentation with plasmid-encoded catalysts leaves the native genome intact and can be transferred across strains, though it depends on sustained co-expression of the folding machinery. Building on the Erv1p–isomerase oxidative folding system described above, we combine CFAI-derived extract preparation, modular extract mixing, and a quantitative Design of Experiments (DoE) framework to independently optimize the chaperone and oxidative folding components of a cell-free reaction [27, 28, 29]. We chose an Erv1p–isomerase oxidative folding strategy rather than a genetically disrupted strain for two reasons: first, we observed that the disruption of reducing pathways in SHuffle is incompatible with the redox-dependent extract activation that drives CFAI performance, resulting in significantly reduced extract activity despite comparable biomass yields. Second, the modular plasmid-based nature of the Erv1p isomerase system could allow deployment across different strains, such as recoded *E. coli* strains offering greater flexibility than genomic knockouts [30, 31]. In strains carrying trxB/gor disruption, including SHuffle, loss of reducing activity can impair ribonucleotide reductase (RNR), which is required for DNA synthesis, and produce a growth defect [20].This defect compromises the scalability and extract quality required by a high-throughput CFAI workflow, making the Erv1p + isomerase strategy a more compatible route to cytoplasmic disulfide bond formation in BL21*DE3 extracts.

Here we move beyond a “one-size-fits-all” approach to extract preparation by establishing a high-throughput, modular CFPS workflow. Cell-Free Autoinduction (CFAI) provides a low-cost, reproducible, and high-yield method for preparing BL21*DE3 extracts [3, 4]. Building on CFAI, we extend this workflow to generate specialized extracts by overexpressing defined chaperone or oxidative-folding systems in the source cells before extract preparation. As with the Colicin M work, these specialized extracts can be combined as interchangeable components for screening.

Complex eukaryotic proteins often require both chaperone assistance and a regulated redox environment to form essential disulfide bonds [15, 16, 26]. To meet both requirements,we expand the mixing strategy to include extracts enriched with oxidative folding enzymes Erv1p with PDI or DsbC. We then validate this approach for the expression of vtPA, the truncated tissue plasminogen activator variant comprising the kringle 2 and protease domains, which together require nine disulfide bonds for activity [32, 33]. Because chaperone and oxidative folding components are supplied as separate, independently tunable extracts, a Design of Experiments (DoE) framework enables systematic optimization of their ratios to maximize soluble yield for each target [27, 28, 29]. Here, we use a quantitative response-surface framework to optimize ratios of chaperone-enriched and oxidative-folding extracts for difficult-to-express targets.

## 2.0 Results and Discussion

The soluble expression of difficult proteins depends on two coupled requirements: a redox environment capable of forming and rearranging native disulfide bonds, and sufficient chaperone capacity to support folding intermediates in solution while redox chemistry proceeds [8, 15, 16]. In a living cell, both are set by host physiology and cannot be adjusted independently. Because cell-free reactions are not constrained by host viability, these requirements can be supplied as separate inputs and dosed against one another.

We tested this idea in stages, first confirming that specialized CFAI extracts retain cell-free productivity (Section 2.1, Figure 1) and then using Colicin M to test whether mixing chaperone-enriched extracts improves soluble expression independent of oxidative folding (Section 2.2). For disulfide-bonded proteins, we found that a genetically oxidizing SHuffle host was incompatible with the CFAI workflow (Section 2.3), which motivated a plasmid-based oxidative folding strategy instead. Combining this strategy with chaperone extracts, we screened folding environments for soluble vtPA (Section 2.4), used Design of Experiments to optimize extract composition and loading (Sections 2.5 and 2.6), and confirmed that the resulting protein is enzymatically active (Section 2.7). Finally, we modeled the materials costs and considered the requirements for scaling the approach (Section 2.8). Together, these experiments yielded soluble and active Colicin M, as well as soluble and enzymatically active vtPA from a common CFAI workflow. By pairing the low modeled materials costs and streamlined preparation of CFAI extracts with modular control of the folding and redox environments, this approach moves cell-free biomanufacturing toward targets that were previously out of reach for the broader cell-free community.

**Figure 1.**
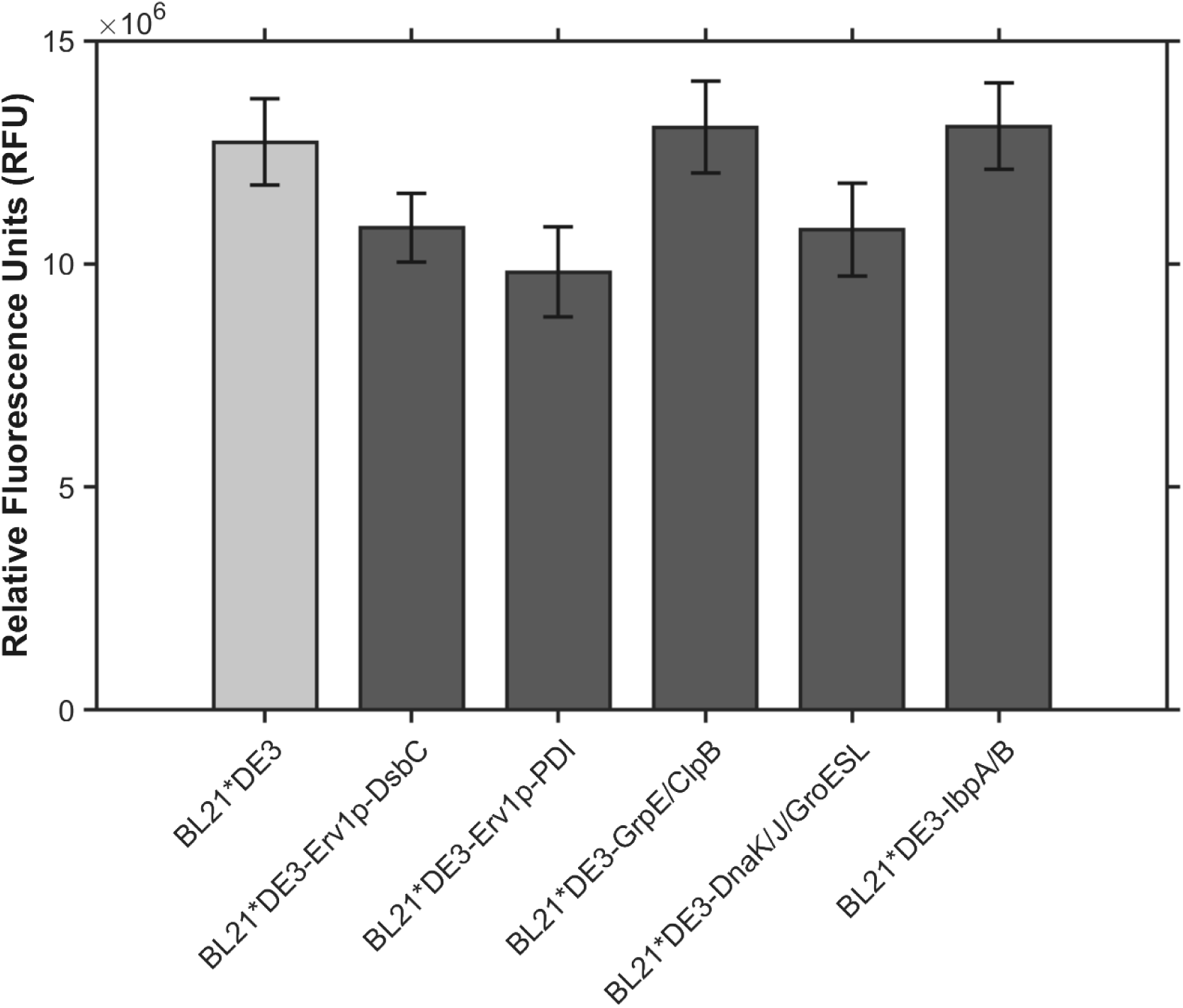
sfGFP expression across BL21*DE3-derived specialized extract strains under CFAI conditions. Relative fluorescence intensity (RFU) of sfGFP produced in cell-free reactions using extracts derived from BL21*DE3 and five specialized BL21*DE3 strains enriched with oxidative folding components (Erv1p–DsbC or Erv1p–PDI) or chaperone systems (GrpE/ClpB, DnaK/J/GroESL, or IbpA/B). All cultures were grown in CFAI media at 30 °C overnight before extract preparation. Error bars represent the standard deviation across biological replicates.

### 2.1 Specialized CFAI Extracts Retain Cell-Free Productivity

To determine whether CFAI could support specialized folding environments without sacrificing extract productivity, sfGFP expression was assessed across five BL21*DE3-derived extracts enriched through cellular overexpression of either oxidative-folding systems (Erv1p–DsbC or Erv1p–PDI) or chaperone systems (GrpE/ClpB, DnaK/J/GroESL, or IbpA/B) (**Figure 1**). All enriched extracts produced sfGFP at levels comparable to the unenriched BL21*DE3 baseline, indicating that overexpression of folding components did not substantially compromise CFAI performance. These results establish that CFAI can generate productive, specialized extracts suitable for modular mixing.

### 2.2 Modular Chaperone Mixing Improves Colicin M Expression

Having established that CFAI can produce functional chaperone-enriched extracts, we first evaluated the chaperone component of the modular platform using Colicin M. Colicin M is an antibacterial bacteriocin produced by *E. coli* that kills susceptible cells by disrupting peptidoglycan biosynthesis, making it a biologically active protein whose successful production can be evaluated through both solubility and antimicrobial function. Because Colicin M is aggregation-prone but lacks disulfide bonds, it also provides a useful model for isolating the effects of chaperone enrichment independently of oxidative folding. We selected three chaperone-enrichment conditions representing complementary stages of proteostasis: GrpE/ClpB, DnaK/J/GroESL, and IbpA/B. Together, these systems encompass ATP-dependent folding and disaggregation as well as small heat-shock-protein-mediated stabilization of aggregation-prone intermediates [6, 7, 9]. Because these plasmids encode multiple folding factors, the resulting extracts represent chaperone-enriched environments rather than isolated individual chaperone activities; consequently, effects cannot be attributed to a single chaperone component.

All chaperone-enriched conditions increased the soluble fraction of Colicin M relative to the unenriched BL21*DE3 control (**Figure 2**). Percent solubility, displayed above each condition in **Figure 2c**, offers a more direct measure of folding efficiency than total protein output, capturing the fraction of expressed protein that avoids aggregation.

**Figure 2.**
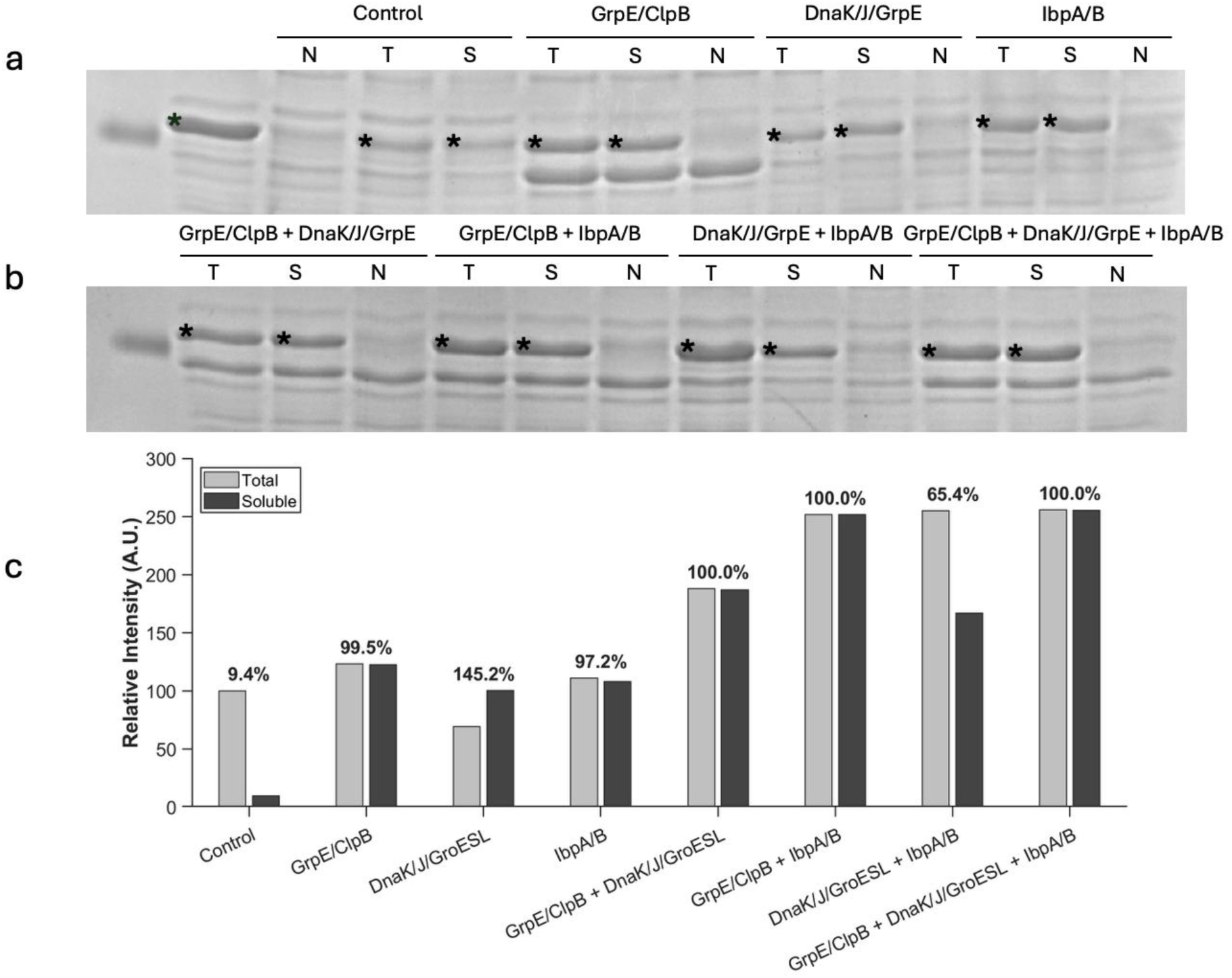
Combinatorial chaperone screening for enhanced soluble Colicin M expression. (a) SDS-PAGE analysis of total and soluble Colicin M, indicated by asterisks (∼29 kDa) produced using unsupplemented BL21*DE3 extract or extracts enriched with GrpE/ClpB (pBB540), DnaK/J/GroESL (pBB550), or IbpA/B (pBB572). A no-DNA extract control was included to identify background bands near the target molecular weight. (b) SDS-PAGE analysis of Colicin M partitioning across pairwise and three-system chaperone combinations. The individual chaperone-enriched extracts shown in (a) were also combined with one another in pairwise and three-way mixtures to test for additive or synergistic effects on solubility. (c) Densitometric quantification of total (gray) and soluble (dark gray) Colicin M fractions, reported as relative integrated density (A.U.) and normalized to the unsupplemented BL21*DE3 extract control. Percent solubility, calculated as the ratio of soluble to total integrated density, is displayed above each condition.

Percent solubility improved markedly across all chaperone-enriched conditions relative to the control, which yielded 9.4% soluble protein (**Figure 2c**). The best conditions reached >95% solubility, a >10-fold improvement, while the weakest supplemented condition still reached 65.4% solubility. Single-system enrichments maintained total protein expression comparable to the control while improving solubility (GrpE/ClpB: 99.5%; IbpA/B: 97.2%), indicating that chaperone supplementation redirects existing expression capacity toward the soluble fraction rather than increasing overall yield. The DnaK/J/GroESL single-system condition showed lower apparent total Colicin M signal than the other individual enrichments. Because sfGFP expression from the same enriched extract remained comparable to the BL21*DE3 baseline (**Figure 1**), this difference is more consistent with variation in gel loading or densitometric recovery than with a generalized loss of extract activity. Among the multi-chaperone combinations, GrpE/ClpB + DnaK/J/GroESL and GrpE/ClpB + IbpA/B both achieved >95% solubility. The combined three-system condition similarly achieved >95% solubility and produced the highest absolute total expression observed across all conditions.

In contrast, the DnaK/J/GroESL + IbpA/B combination yielded 65.4% solubility despite both systems performing well independently. This pattern is consistent with the known cooperative roles of these chaperone systems, although the densitometry data alone do not establish causation [6, 8]. The recovery of >95% solubility in the combined three-system condition further supports a cooperative dynamic. One possible explanation for this pattern is the composition of the pBB550 extract, which overexpresses DnaK and DnaJ but does not provide a corresponding increase in GrpE, the nucleotide exchange factor required for efficient cycling of the DnaK system. Combining pBB550 with pBB540 provides additional GrpE as well as the ClpB disaggregate, which may improve substrate release and recycling during folding. A similar effect may contribute to the lower solubility observed when the pBB550 and IbpA/B-enriched extracts were combined, as proteins stabilized by small heat-shock proteins can require downstream DnaK- and ClpB-mediated processing for productive refolding [6, 8].

To confirm that the solubility improvements translated to a biologically active product, we performed a cell-kill assay against E. coli K-12. Cell-free–expressed Colicin M exhibited clear antimicrobial activity, with increasing extract volumes producing greater reductions in bacterial survival. Control cells and cells treated with negative-control CFPS extract exhibited confluent growth, demonstrating that the CFPS reaction components themselves do not inhibit bacterial growth. Similarly, cells treated with 2.5 µL Colicin M extract also exhibited confluent growth. In contrast, treatment with 7.5 µL Colicin M extract substantially reduced colony formation, while 12.5 µL extract resulted in near-complete elimination of viable colonies. Representative plates are shown in **Figure S1**, and quantitative colony counts are summarized in **Table 1**. These results confirm that the modular chaperone-mixing strategy yields soluble and functionally active protein, validating the approach and establishing a benchmark before advancing disulfide-containing targets.

**Table 1.** Inhibitory Activity of Cell-Free Expressed Colicin M against *E. coli* K-12.

| Colicin M Extract (μL) | Colony Forming Units (CFU) |
| --- | --- |
| Cells only (0 μL CFPS) | Lawn |
| 12.5 negative-control CFPS | Lawn |
| 2.5 | Lawn |
| 7.5 | 1730 ± 360 |
| 12.5 | 3.3 ± 1.5 |

### 2.3 SHuffle T7 Is Incompatible with the CFAI Workflow, Motivating a Plasmid-Based Oxidative Folding Strategy

To test whether a genetically oxidizing host could extend CFAI to proteins requiring disulfide bonds, we compared extract productivity from BL21*DE3 and SHuffle T7 grown under conventional 2xYTPG and CFAI conditions. Under conventional conditions, both strains reached comparable OD600 values and yielded similarly productive extracts, confirming that SHuffle T7 can support cell-free protein synthesis when grown conventionally (**Figure 3a, b**). Under CFAI conditions, however, their extract performance diverged. Although SHuffle T7 again reached an OD600 comparable to that of BL21*DE3 (**Figure 3c**), extracts prepared from CFAI-grown SHuffle T7 exhibited markedly reduced sfGFP expression (**Figure 3d**). Because the *ΔtrxB* and *Δgor* knockouts that establish SHuffle’s oxidative cytoplasm also alter redox metabolism more broadly, this loss of activity may reflect disruption of the redox-dependent extract activation mechanism underlying CFAI performance [4].

**Figure 3.**
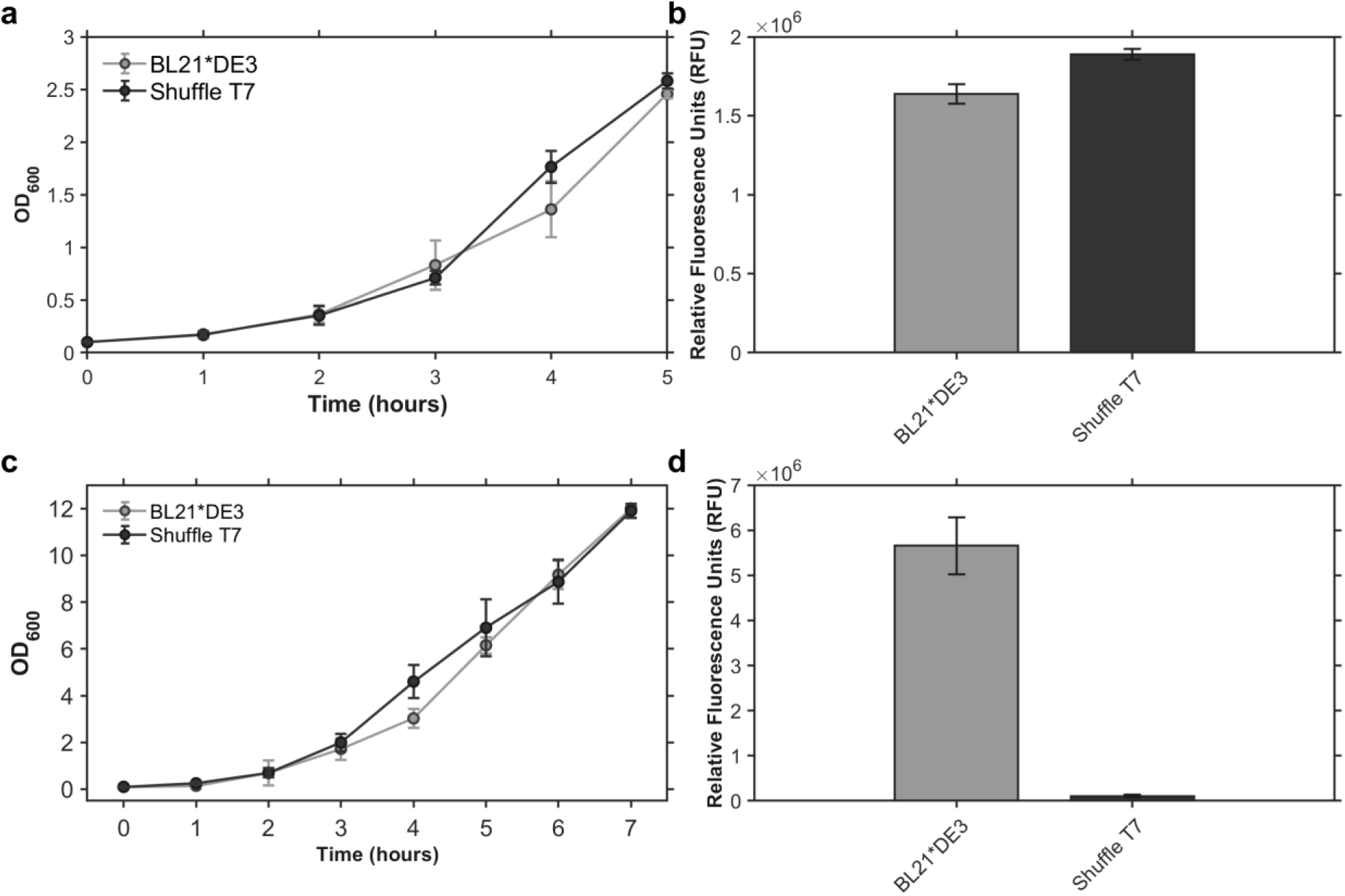
Comparison of BL21*DE3 and SHuffle T7 extract productivity under standard and CFAI growth conditions. (a) Growth curves of BL21*DE3 and SHuffle T7 in standard 2xYTPG media showing comparable OD600 profiles over five hours. (b) sfGFP expression from extracts derived from 2xYTPG-grown cultures, demonstrating that both strains produce functional extracts under standard conditions. (c) Growth curves of BL21*DE3 and SHuffle T7 under CFAI conditions, where both strains again reach comparable OD600 values. (d) sfGFP expression from CFAI-derived extracts showing a marked reduction in SHuffle T7 productivity relative to BL21*DE3 despite equivalent biomass yields. All sfGFP expression values are reported as relative fluorescence units (RFU). Error bars represent standard deviation across biological replicates.

Although SHuffle T7 can produce functional extracts following conventional growth, it could not provide productive oxidizing extracts under the CFAI conditions tested (**Figure 3**). We therefore retained BL21*DE3 as the CFAI host and instead expanded the range of protein classes accessible to the CFAI platform by introducing oxidative folding capacity through plasmid - encoded disulfide bond–forming systems. Specifically, the expression of Erv1p, a sulfhydryl oxidase that catalyzes the oxidation of cysteine thiols to initiate disulfide bond formation, was paired with either protein disulfide isomerase (PDI) or DsbC. PDI and DsbC function as disulfide bond isomerases, facilitating the rearrangement and correction of non-native disulfide bonds and thereby complementing Erv1p-mediated oxidation in modular CFAI extracts.

### 2.4 Screening Modular Folding Environments for Soluble vtPA Expression

Having validated the chaperone mixing strategy on an aggregation-prone substrate, we next assessed whether the platform could accommodate a more demanding folding challenge: a truncated variant of human tissue plasminogen activator (vtPA) comprising the kringle 2 and protease domains, which together require nine disulfide bonds for proper structure and function [33]. Unlike Colicin M, vtPA presents a dual hurdle: it requires both chaperone-assisted prevention of aggregation and a regulated oxidative environment to form and isomerize its disulfide bonds, making it a rigorous benchmark for testing the full capabilities of the modular mixing platform [15, 16].

To identify extract conditions capable of supporting soluble vtPA expression, we performed a preliminary screen across a 2 × 3 matrix of BL21*DE3-derived extract environments, crossing two oxidative folding systems (Erv1p–DsbC or Erv1p–PDI) against three chaperone backgrounds (GrpE/ClpB, DnaK/J/GroESL, or IbpA/B). Each combination was evaluated under both reducing and non-reducing conditions, with expression monitored by FluoroTect green-lysine labeling.

Among the six extract combinations tested, DnaK/J/GroESL supplementation consistently produced the strongest soluble vtPA expression for both oxidative folding systems (**Figure 4**). We therefore carried both Erv1p–DsbC and Erv1p–PDI forward into DoE optimization, each paired with DnaK/J/GroESL, to evaluate the two isomerases under the lead chaperone background.

**Figure 4.**
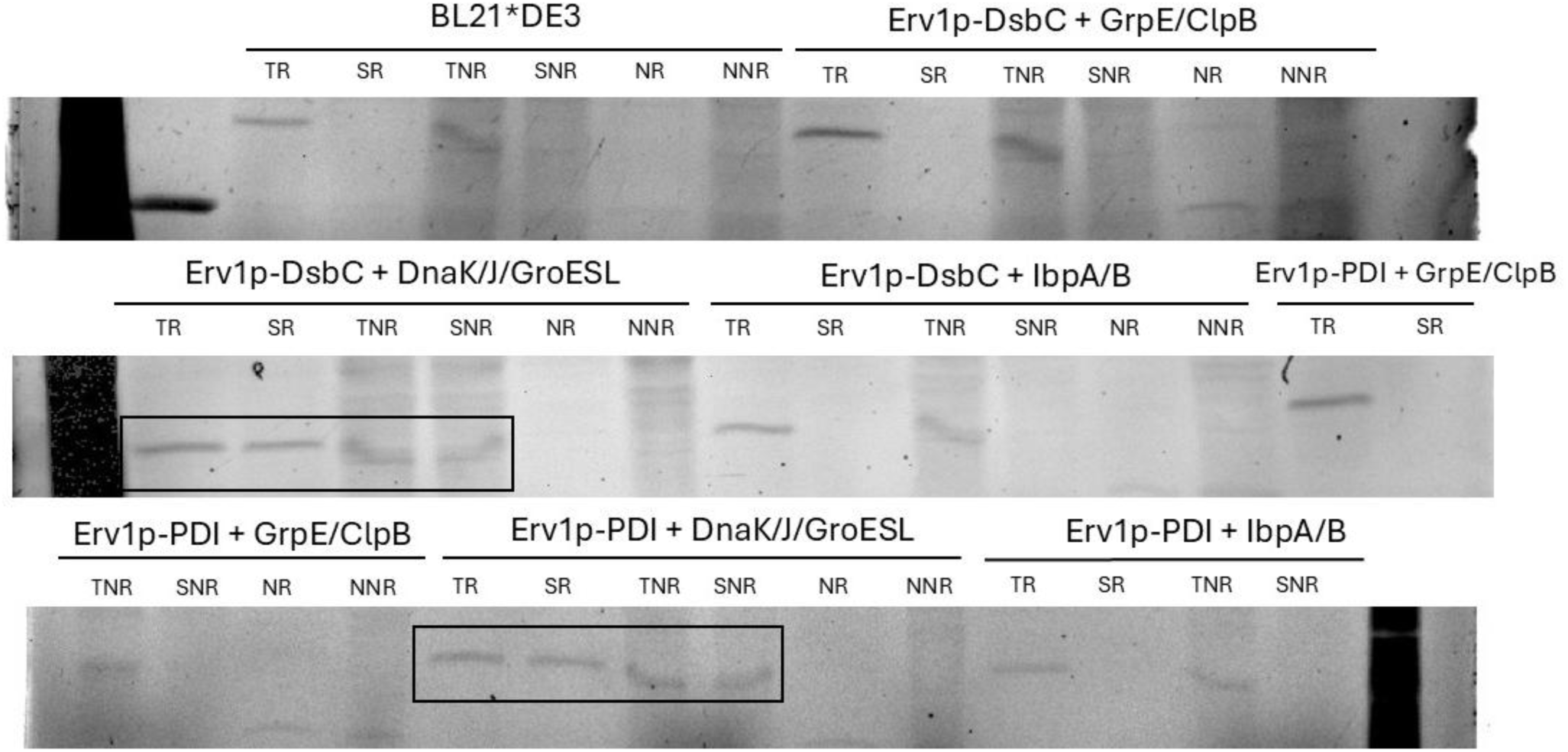
Preliminary screen of oxidative folding and chaperone extract combinations for soluble vtPA expression. FluoroTect-labeled SDS-PAGE gels showing vtPA expression across a 2 × 3 matrix of oxidative folding–chaperone combinations: Erv1p–DsbC or Erv1p–PDI, each paired with GrpE/ClpB (pBB540), DnaK/J/GroESL (pBB550), and IbpA/B (pBB572). A non-supplemented BL21*DE3 extract is included as a baseline. For each condition, lanes are shown in the following order: total reducing (TR), soluble reducing (SR), total non-reducing (TNR), soluble non-reducing (SNR), negative reducing (NR), and negative non-reducing (NNR). Green boxes denote the lead DnaK/J/GroESL condition, which produced the strongest soluble vtPA signal for both oxidative folding systems.

The robust performance of both oxidative folding systems when paired with DnaK/J/GroESL highlights the importance of appropriate chaperone supplementation. As a native prokaryotic isomerase, DsbC is optimized to function in bacterial folding environments with chaperones such as DnaK. DsbC also possesses intrinsic chaperone activity in addition to its isomerase function, allowing it to shield hydrophobic patches on the nascent vtPA chain while correcting non-native disulfide bonds. This dual role may be critical for a substrate with nine disulfide bonds [15, 16]. PDI similarly demonstrated effective performance when supplied with bacterial chaperones, suggesting that eukaryotic isomerases can function in prokaryotic systems when paired with compatible folding partners. These findings indicate that successful expression of multi-disulfide substrates requires careful matching of isomerase and chaperone components, and this matching is examined quantitatively in the DoE optimization described in Section 2.5.

### 2.5 Establishment of DoE Optimization Framework

With the DnaK/J/GroESL chaperone system identified as the lead configuration for vtPA expression, we transitioned to a systematic Design of Experiments (DoE) framework to quantitatively map the relationship between extract composition and soluble vtPA yield. While the preliminary screen identified DnaK/J/GroESL as the lead chaperone background in combination with either Erv1p–DsbC or Erv1p–PDI, it could not determine the optimal concentrations or volumetric ratios required to maximize folding efficiency, a question that a factorial design is well suited to address.

We developed a full factorial design to investigate two critical parameters: total translational capacity (extract loading) and the balance between chaperone and oxidative folding system activity. As detailed in **Table 2**, the design space explored extract loadings from 25% to 40% (v/v) and chaperone ratios ranging from 0% to >95%. We applied this same design space to both the Erv1p–DsbC and Erv1p–PDI oxidative folding systems, each evaluated against the DnaK/J/GroESL chaperone background, to determine whether the optimal extract composition is conserved between them. To ensure the stability of cell-free reactions across the experimental timeline, independent replicate runs, including center-point replicates, were performed for each oxidative folding system. These replicates served as internal controls to verify the reproducibility of the synthesis environment, allowing for a robust characterization of the interaction between translational capacity and chaperone availability for the 9-disulfide vtPA protease domain.

**Table 2.** Design of Experiments (DoE) Matrix for Optimization of vtPA Soluble Expression.

| Run | Total<br>Extract % | Ratio (Erv1p–<br>DsbC or Erv1p–<br>PDI :<br>DnaK/J/GroESL) | Condition<br>Type |
| --- | --- | --- | --- |
| 1 | 25% | 0 / 3c | Control |
| 2 | 25% | 25 / 75 | Experimental |
| 3 | 25% | 50 / 50 | Experimental |
| 4 | 25% | 75 / 25 | Experimental |
| 5 | 25% | 100 / 0 | Control |
| 6 | 33% | 0 / 100 | Control |
| 7 | 33% | 25 / 75 | Experimental |
| 8 | 33% | 50 / 50 | Center point |
| 9 | 33% | 75 / 25 | Experimental |
| 10 | 33% | 100 / 0 | Control |
| 11 | 40% | 0 / 100 | Control |
| 12 | 40% | 25 / 75 | Experimental |
| 13 | 40% | 50 / 50 | Experimental |
| 14 | 40% | 75 / 25 | Experimental |
| 15 | 40% | 100 / 0 | Control |
| 16 | 33% | 50 / 50 | Replicate |
| 17 | 33% | 50 / 50 | Replicate |
| 18 | 33% | 0 / 100 | Replicate |

### 2.6 Identification of Robust Conditions for vtPA

Densitometric analysis across the 18 DoE conditions demonstrated that soluble vtPA yield depends on both extract loading and the chaperone-to-isomerase ratio for both the Erv1p–DsbC and Erv1p–PDI systems (**Figure 5**). For each oxidative folding system, values were normalized to its respective center-point replicate (Condition 8; 33% total extract, 50:50 oxidative folding system: DnaK/J/GroESL), assigned a reference value of 100 A.U. The full design matrix with coded factor levels and dthe resulting normalized responses for both systems is given in **Table S1**. All reactions were performed under reducing conditions to isolate the effects of extract composition and chaperone ratio, ensuring that observed differences in solubility reflect folding support rather than variations in redox environment.

**Figure 5.**
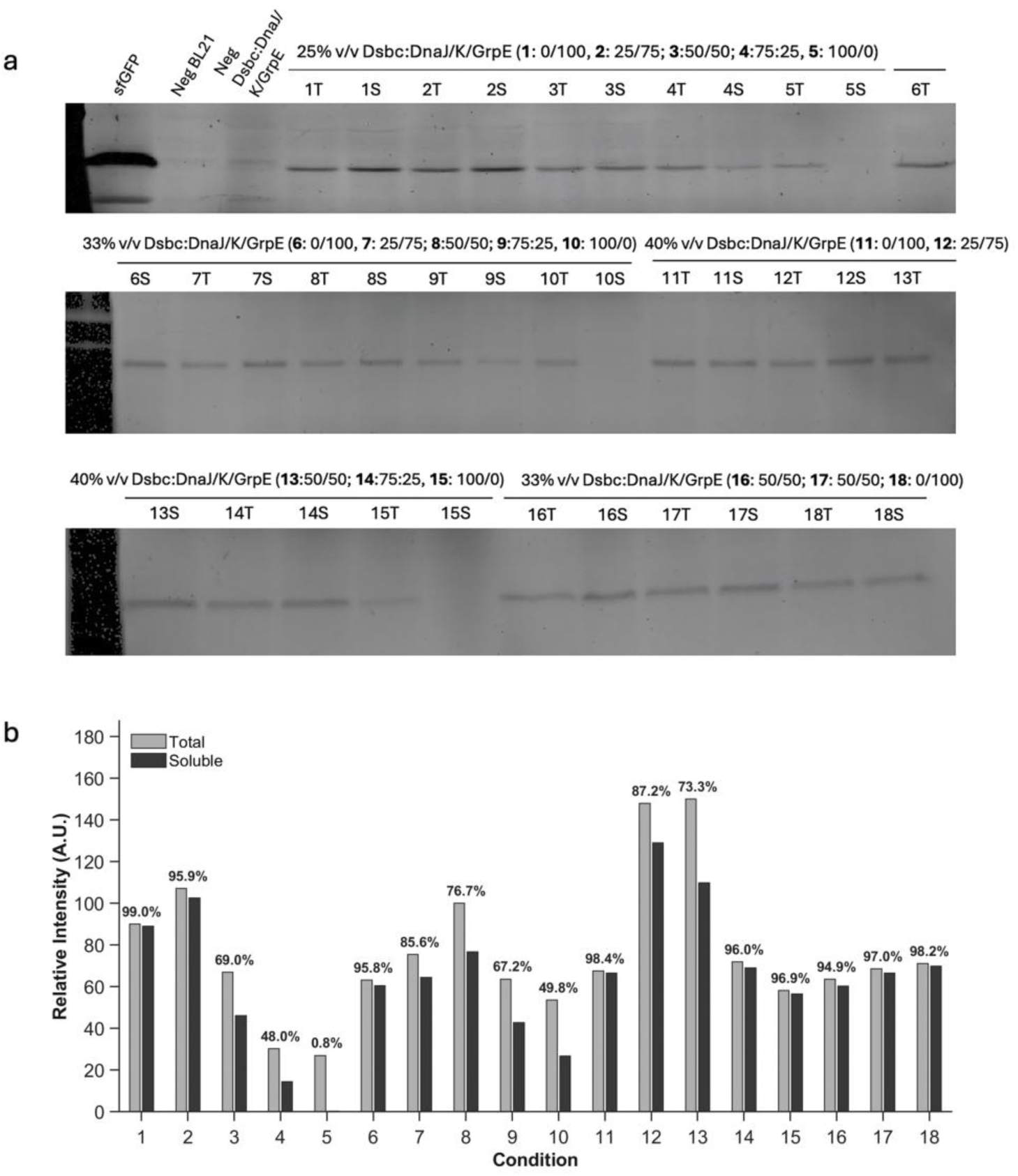
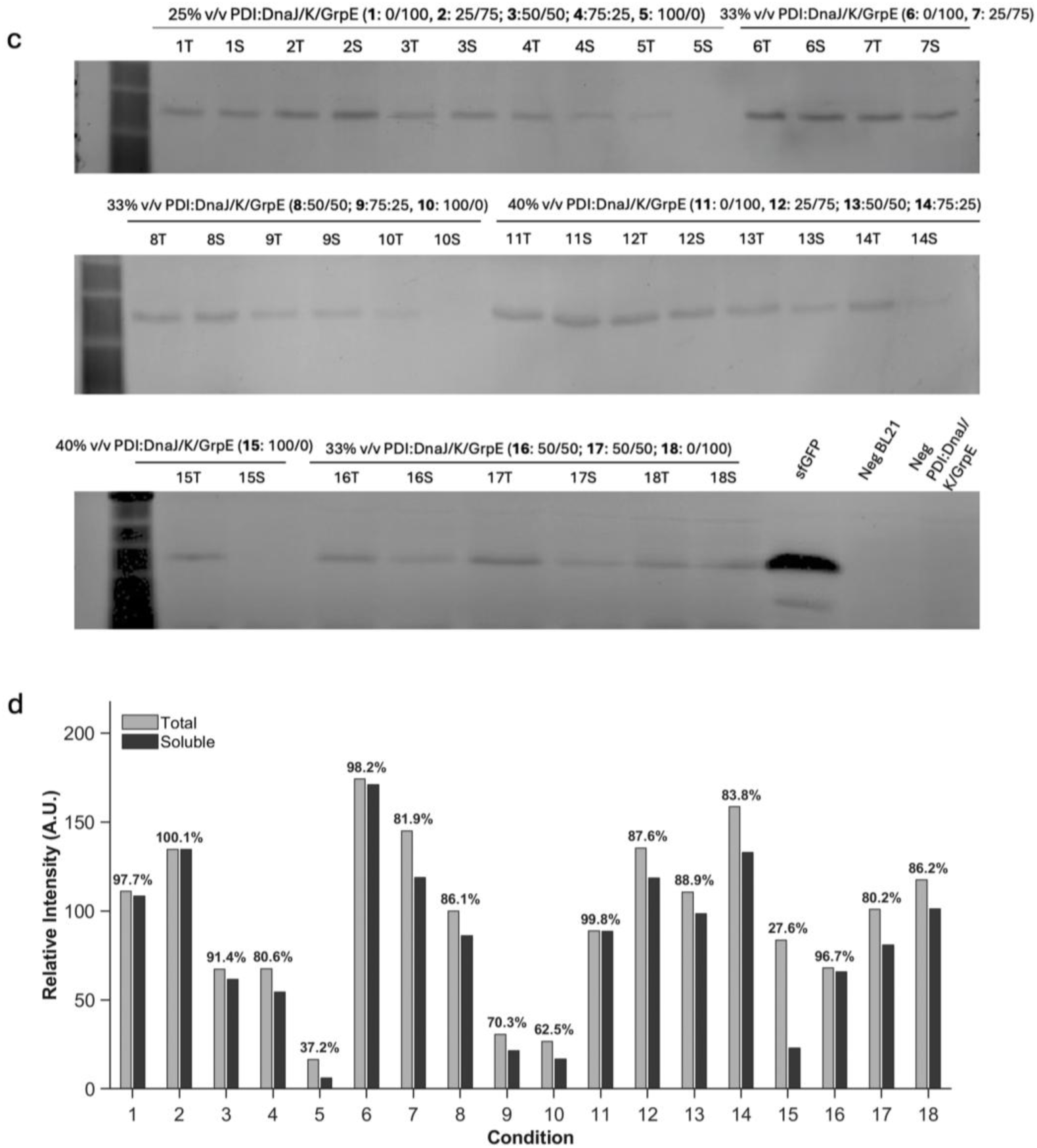
DoE optimization of soluble vtPA expression across extract loading and chaperone-to-isomerase ratio for the Erv1p–DsbC and Erv1p–PDI systems. (a) SDS-PAGE gels showing total (T) and soluble (S) protein fractions for all 18 DoE conditions of the Erv1p–DsbC system. Conditions 1–15 represent unique DoE design points; Conditions 16 and 17 are center-point replicates (33% total extract, 50:50 ratio), and Condition 18 is a replicate of the 0:100 (chaperone-only) condition, each performed as a wholly separate reaction to assess experimental reproducibility. (b) Densitometric quantification of total (grey) and soluble (green) fractions for the Erv1p–DsbC system, expressed as relative integrated density (A.U.) and normalized to Condition 8. (c) Corresponding SDS-PAGE gels for all 18 DoE conditions of the Erv1p–PDI system, following the same condition layout. (d) Densitometric quantification of total (grey) and soluble (green) fractions for the Erv1p–PDI system, normalized to its respective Condition 8. In (b) and (d), percent solubility values, calculated as the ratio of soluble to total integrated density, are displayed above each condition. Conditions are numbered according to the DoE matrix in Table 2.

For the Erv1p–DsbC system (**Figure 5a, b**), densitometric analysis revealed a clear threshold effect across the design space. Solubility remained high when DnaK/J/GroESL-supplemented extracts constituted the majority of the extract composition, but declined sharply once the Erv1p– DsbC extract exceeded 50% of the total extract volume, a pattern that held across all three loading levels. Conditions in which the Erv1p–DsbC extract constituted 75% or more of the total extract consistently produced the lowest soluble fractions regardless of total extract concentration, pointing to excess isomerase enrichment rather than loading level as the primary driver of aggregation. The poorest-performing conditions dropped as low as 0.8% solubility, reinforcing that Erv1p-DsbC dominant compositions are poorly tolerated. High Erv1p-DsbC loading did not simply fail outright, however: Conditions 12 and 13 produced the highest absolute total expression across the Erv1p-DsbC design space, yet their solubility remained comparatively modest (87.2% and 73.3%), indicating that elevated total yield at high Erv1p-DsbC loading does not translate proportionally into soluble protein. By contrast, chaperone-dominant and balanced conditions at higher loading consistently achieved both strong expression and high solubility, exceeding 95% in several cases, identifying chaperone-rich compositions as the most favorable for soluble vtPA expression. The center-point replicates returned solubilities of 94.9–98.2%, demonstrating good reproducibility at the center of the experimental design space.

Applying the identical DoE framework to the Erv1p–PDI system (**Figure 5c, d**) revealed a broadly consistent dependence on chaperone availability, with several informative differences. As with the Erv1p-DsbC system, the strongest soluble yields were associated with chaperone-rich compositions, several of which exceeded 98% solubility alongside strong absolute expression. Erv1p-PDI dominant conditions again produced the weakest performance, with isomerase-heavy conditions (>95%) dropping as low as 27.6% solubility, consistent with the isomerase-excess aggregation observed for Erv1p-DsbC. Intermediate chaperone-to-isomerase ratios at higher loading maintained robust expression with solubilities in the mid-80% range, indicating that the Erv1p-PDI system tolerated a balanced folding environment well, provided chaperone support remained substantial.

To quantify the contribution of each factor and identify the optimal extract composition, a second-order response-surface model was fit to the normalized soluble vtPA signal for each oxidative folding system, incorporating main effects for extract loading and chaperone ratio, their interaction, and quadratic terms for both factors (**Figure 6**). Soluble integrated density was used as the response variable rather than percent solubility, as the latter saturates near complete solubilization and does not capture absolute yield. For the Erv1p-DsbC system, the model accounted for the majority of the observed variation (R² = 0.80, adjusted R² = 0.72; overall model p < 0.001). Chaperone ratio was the dominant factor (p < 0.001), with extract loading (p = 0.004) and a significant loading by ratio interaction (p = 0.029) also contributing, indicating that the effect of extract loading on soluble yield depends on the chaperone ratio. Significant negative curvature in the chaperone ratio term (p = 0.017) further supports the presence of an internal optimum in chaperone fraction rather than a monotonic response.

**Figure 6.**
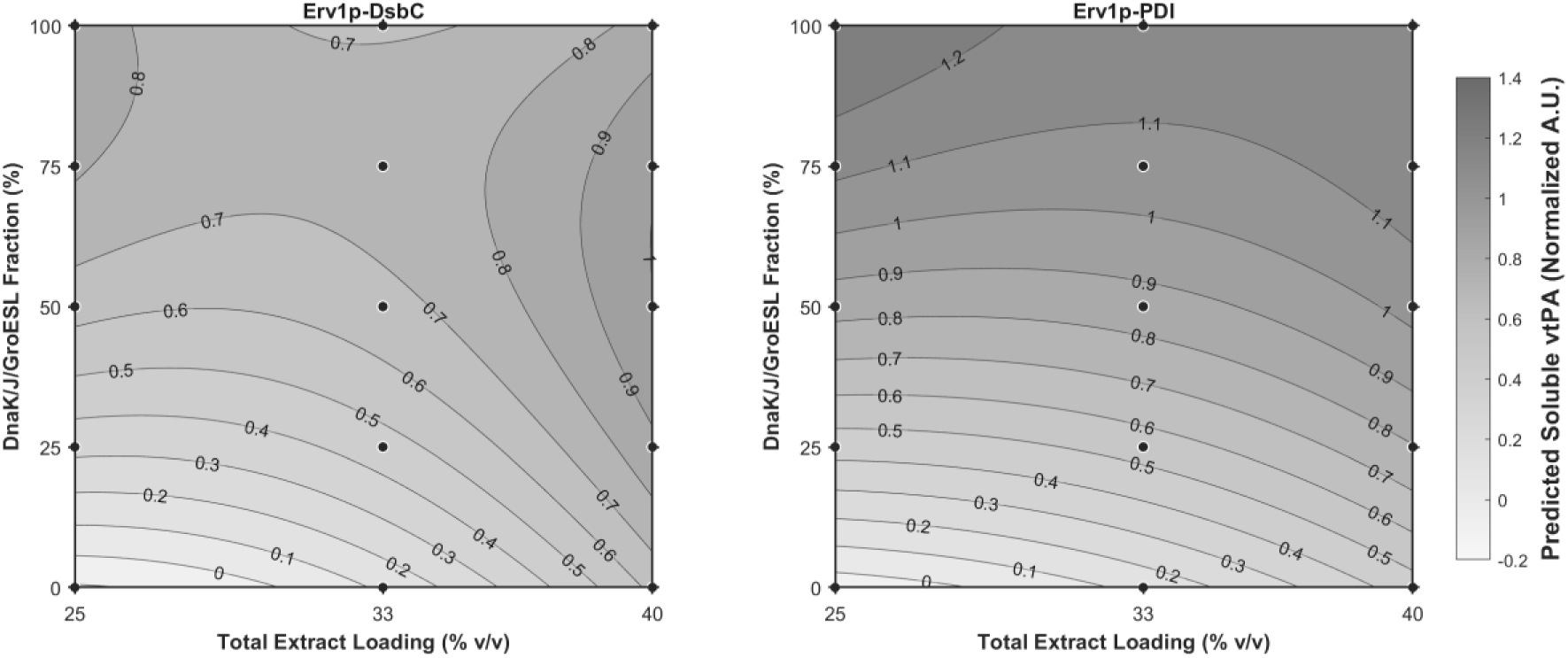
Response-surface models of the soluble vtPA yield as a function of total extract loading and DnaK/J/GroESL chaperone fraction for the Erv1p-DsbC (left) and Erv1p-PDI (right) systems. Filled contours represent the predicted normalized soluble vtPA signal (A.U) from a second order least squares fit; black markers denote the 18 experimental design points from the DoE matrix in Table 2. Both panels share a single colour scale spanning the pooled range of the two fitted surfaces (-0.2 to 1.4 A.U.), so shading is directly comparable between panels. Slightly negative predicted values near the low-loading, low-chaperone corner reflect the empirical nature of the quadratic approximation and are not physically meaningful. The Erv1p-DsbC system exhibits a diagonal ridge along which the chaperone fraction giving maximal predicted yield decreases as extract loading increases, reflecting a significant loading-by-ratio interaction, whereas the Erv1p-PDI system shows near horizontal contours consistent with chaperone ratio as the only resolved factor.

For the Erv1p-PDI system, the model similarly captured the observed variation (R² = 0.70, adjusted R² = 0.57; overall model p < 0.01), but chaperone ratio was the sole significant factor (p < 0.001); neither extract loading, the interaction, nor the quadratic terms reached significance. Soluble vtPA yield in the Erv1p-PDI system is therefore governed almost exclusively by chaperone availability, with no resolvable dependence on total extract loading across the tested range. Together, these models identify chaperone fraction as the dominant predictor of soluble vtPA yield in both oxidative folding systems.

Model adequacy was assessed against pure error estimated from the replicated runs. Lack of fit was not significant at the 0.05 level for either system (Erv1p-DsbC, F(9,3) = 6.11, p = 0.082; Erv1p-PDI, F(9,3) = 1.04, p = 0.548), and both models exceeded the conventional adequate-precision threshold of 4 for resolving the response across the design space (11.2 and 7.8, respectively). Because pure error is estimated from only three degrees of freedom, however, these tests have limited power and are interpreted as an absence of evidence against the second - order models rather than confirmation of their adequacy. For both systems the predicted R² (0.49 for Erv1p-DsbC, 0.20 for Erv1p-PDI) differs from the adjusted R² by more than 0.20, indicating limited predictive capability beyond the measured design points; the fitted surfaces are therefore interpreted as descriptive of trends across the tested range rather than as predictive mod els for extrapolation. Full analysis of variance and coefficient estimates for both systems are provided in **Tables S2** and **S3**.

Although both oxidative folding systems followed similar overall trends, the response-surface models revealed an important difference between them. The Erv1p–DsbC system exhibited a significant interaction between extract loading and chaperone ratio, whereas the Erv1p–PDI system was driven primarily by chaperone ratio alone. These results suggest that optimization of the DsbC system requires balancing translational capacity and folding support. For the PDI system, no effect of total extract concentration was resolved across the range tested; given the higher residual variability of the PDI model, this reflects lack of detectable dependence rather than demonstrated insensitivity.

The modular mixing strategy enabled rapid evaluation of both oxidative folding systems across a broad design space without requiring strain re-engineering or new extract preparations for each condition. By treating specialized extracts as interchangeable components, the full DoE matrix could be screened within a single workflow, demonstrating the utility of modular extract mixing for optimization of complex cell-free environments [10, 11, 12].

### 2.7 Functional Validation of vtPA Activity

To confirm that the solubility gains observed under the DoE conditions translated to a biologically active product, we performed a fluorescence-based protease activity assay using 50 μM AMC-conjugated substrate with soluble fractions from three optimized extract systems: >95% DnaK/J/GroESL extract, a 50:50 Erv1p–DsbC:DnaK/DnaJ/GroESL extract mixture, and a 50:50 Erv1p–PDI:DnaK/DnaJ/GroESL extract mixture, each prepared at 40% total extract loading. As shown in Figure 7a, all three conditions exhibited increasing fluorescence over time, indicating successful substrate cleavage by cell-free–expressed vtPA. However, the Erv1p– DsbC:DnaK/DnaJ/GroESL condition produced a markedly greater fluorescence increase than the other two systems over the course of the assay. Quantification of the kinetic traces further supported this trend (**Figure 7b**), with the Erv1p–DsbC:DnaK/DnaJ/GroESL condition exhibiting the highest specific activity (2775 ± 292 RFU/min), compared to 1840 ± 790 RFU/min for the DnaK/J/GroESL-only condition and 1518 ± 380 RFU/min for the Erv1p– PDI:DnaK/DnaJ/GroESL condition.

**Figure 7.**
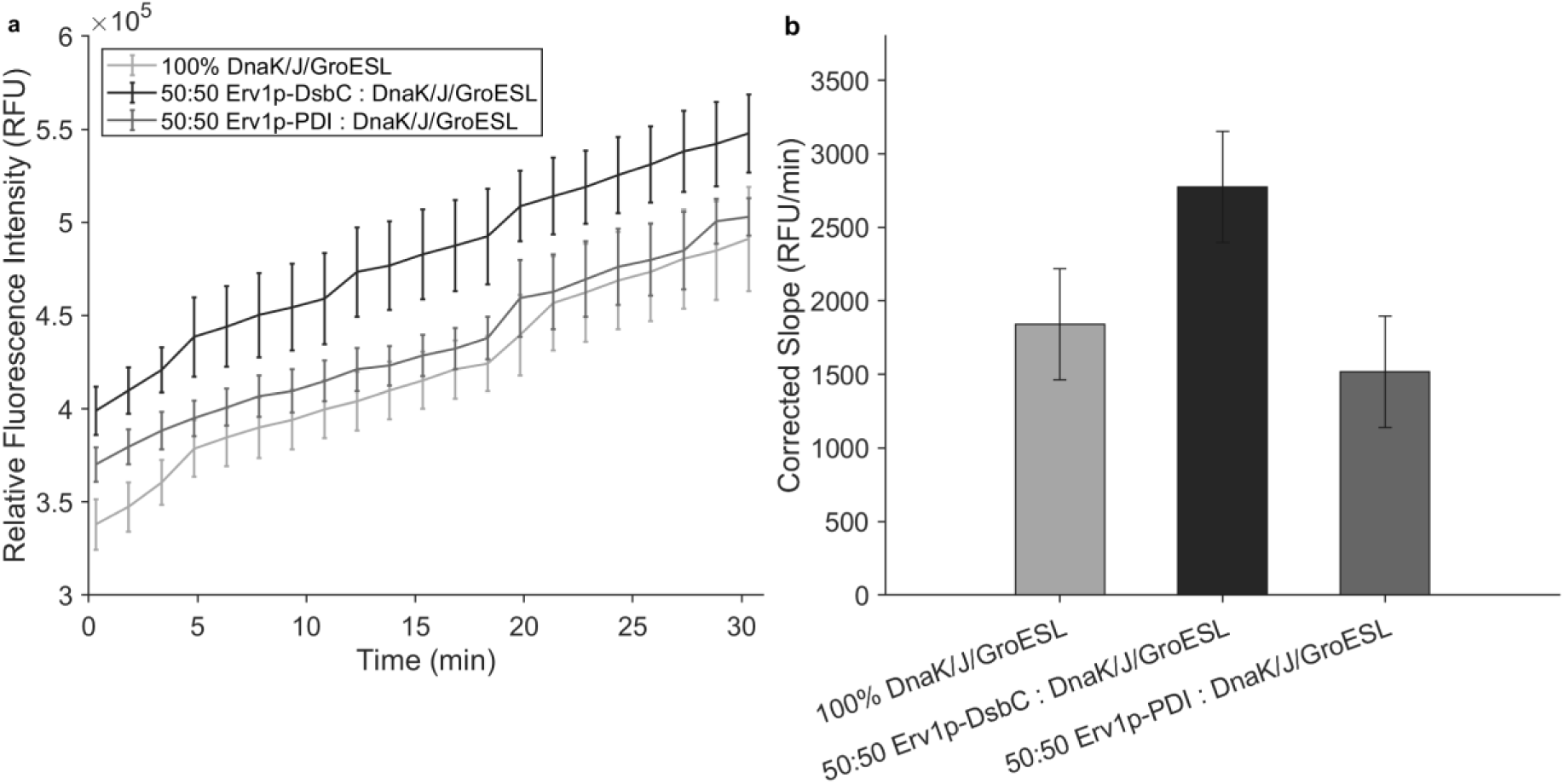
Functional validation of cell-free expressed vtPA across optimized extract systems. (a) Fluorogenic protease activity assay of vtPA produced using 40% total extract loading with either >95% DnaK/J/GroESL extract (Condition 11), a 50:50 Erv1p–DsbC:DnaK/DnaJ/GroESL extract mixture (Condition 13 of the Erv1p-DsbC design), or a 50:50 Erv1p–PDI:DnaK/DnaJ/GroESL extract mixture (Condition 13 of the Erv1p-PDI design). Soluble fractions were diluted 5× and incubated with 50 μM D-Ile-Pro-Arg-AMC substrate. Protease activity was monitored by AMC fluorescence release following substrate cleavage. Fluorescence values were corrected by subtracting both a negative CFPS control and substrate background. Error bars represent SEM (n = 3). (b) Corrected vtPA-specific activity calculated from the kinetic traces shown in panel (a). Values above bars indicate mean corrected slope ± propagated SEM.

Because a purified vtPA standard was not included in this study, activity is reported as relative fluorescence (RFU/min) rather than absolute specific activity, an acceptable practice in the field. Future studies will benchmark the optimized cell-free product against purified vtPA standards to enable direct comparison with native enzyme activity.

The observed activity trends are consistent with the expected roles of the folding systems tested. While DnaK/J/GroESL functions as a general folding chaperone that suppresses aggregation, it does not directly catalyze disulfide bond formation. Consequently, although the DnaK/J/GroESL-only condition produced measurable vtPA activity, its performance remained lower than that of the Erv1p-DsbC condition, suggesting that productive disulfide bond formation remains an important determinant of vtPA function. The superior performance of the Erv1p–DsbC system relative to Erv1p–PDI indicates that DsbC may be more suitable for vtPA folding in the CFAI-based cell-free environment [15]. As a native bacterial disulfide isomerase, DsbC may function more efficiently within the biochemical environment of *E. coli* extracts, enabling more effective correction of non-native disulfide bonds during vtPA folding specifically.

Although both oxidative folding systems yielded highly soluble vtPA, only the Erv1p–DsbC condition produced a substantial gain in activity relative to the chaperone-only condition. This suggests that solubility alone is not a reliable indicator of correct folding and that the Erv1p– DsbC system more effectively promotes formation of the active vtPA conformation under these conditions. The measurable activity observed in the chaperone-only condition further indicates that some productive folding can occur in the absence of an engineered oxidative folding system, although the Erv1p–DsbC system is required to maximize functional output.

These results provide the first functional validation of the modular extract-mixing strategy for vtPA, a disulfide-rich therapeutic target, and demonstrate that the choice of oxidative folding catalyst can meaningfully influence the functional output of a complex disulfide bonded protein, not just its solubility. By establishing the 50:50 Erv1p–DsbC + DnaK/J/GroESL composition at 40% extract loading as the lead condition for active vtPA production, this work shows that the modular platform can be tuned to identify protein-specific folding solutions, providing a generalizable framework for optimizing the cell-free synthesis of other therapeutically relevant disulfide bonded proteins.

### 2.8 Cost and Scalability of Modular Extract Mixing

We evaluated the economics of modular extract mixing with an ingredient-based model of known materials consumed per 15 µL reaction, separating reaction reagents, extract preparation, and purchased folding supplements. This framework complements prior CFPS cost analyses and supplementation strategies [13, 14, 34, 35]. CFAI and 2xYTPG extract costs were modeled using their respective preparation workflows [3, 5, 36, 37]. These are materials-consumption estimates rather than complete experimental expenditures. Target-template DNA, selection-plate materials, water, titration and freezing materials remain unpriced, while analytical labeling, activity assays, consumables, labor, and equipment are excluded. The model uses nominal reagent concentrations; unresolved differences between nominal and weighed stock formulations also preclude an exact accounting of experimental spending.

CFAI avoids IPTG induction and the associated monitoring and timed intervention [4, 5]. In the updated model, CFAI growth medium costs $13.06/L, whereas 2xYTPG medium costs $10.61/L before addition of $11.34/L IPTG at the modeled 1 mM dose, giving $21.95/L for medium plus inducer. With buffer and stock-preparation allocations included, standard CFAI and 2xYTPG extracts cost approximately $1.66 and $5.88 per mL of usable extract, respectively. This approximately 72% reduction depends on assumed usable yields of 10 mL/L culture for CFAI and 4.29 mL/L for 2xYTPG. The CFAI value is a modeling assumption, and equal usable yields among enriched strains have not been established experimentally.

The five folding-system plasmids used here—pBB540, pBB550, pBB572, pCOLA-Gent-EM7-Erv1p–DsbC, and pCOLA-Gent-EM7-Erv1p–PDI—are assigned an acquisition cost of $94 each, or $470 for the set [38]. The model allocates one construct and $27.80 in competent cells across 10 L of culture per enriched strain stock. Acquisition is therefore included in the extract unit cost rather than treated as a zero-cost input or added again to each reaction. The resulting enriched - extract estimates are $2.60/mL for DTT-containing chaperone extracts and $2.54/mL for DTT-free extracts. Mixtures of separately enriched extracts are costed by their volume fractions. Enrichment avoids a purchased-protein supplement charge, but its materials cost still depends on stock reuse, extract recovery, and expression burden.

At the standard 5 µL extract loading, the nominal PANOx reaction reagents contribute $0.1789 per 15 µL reaction before unpriced DNA. Adding extract gives known-material subtotals of $0.1872 for standard CFAI, $0.2083 for standard 2xYTPG, $0.1919 for CFAI with DTT-containing chaperone extract, and $0.1916 for DTT-free CFAI with oxidative machinery or both folding systems. For the lead vtPA condition, 40% extract corresponds to 6 µL per reaction and raises the CFAI subtotal to $0.1942. All vtPA extracts were prepared without DTT (Section 4.2). Under the equal-yield and stock-allocation assumptions, changing the ratio of these enriched extracts at fixed loading does not change their modeled unit cost, whereas changing the total extract loading does.

Purchased supplementation was modeled using GeneFrontier DnaK Mix, GroE Mix, and DsbC Set at their label doses, normalized to 15 µL final reaction volume [39]. At the selected 50 mL-capacity package tier, DnaK Mix and GroE Mix together add $6.5556 per reaction, and DsbC Set adds $1.8354, giving $8.3910 for both supplement classes. With 5 µL of 2xYTPG extract, the corresponding known-material subtotals are $6.7639 for chaperones, $2.0434 for the oxidative set, and $8.5990 for both. The standard-loading comparison is summarized in Table S4 and Figure S2. At the 6 µL loading used for the lead vtPA comparison, both purchased sets give $8.6048 per reaction versus $0.1942 for CFAI mixing, a 44.3-fold difference or 97.7% lower modeled known-material cost. This comparison represents reagent purchasing scenarios, not experimentally matched protein output.

Package size and supplier choice materially affect the comparison. At 6 µL extract loading, the combined GeneFrontier scenario ranges from $8.6048 per reaction using 50 mL-capacity packages to $10.2188 using 10 mL packages and $12.7238 using 0.5 mL packages [39]. These unit costs assume consumption at the selected package price; purchasing whole packages from zero inventory is a separate expense. Other oxidative-support options in the model are the NEB Disulfide Bond Enhancer at $3.60 per 15 µL reaction [40] and ProSpec DsbC alone at $2.85 per reaction from a 1 mg package at an illustrative 100 µg/mL dose [41]. These are supplement-only prices, not complete CFPS reaction costs. The Ginkgo DSB Booster estimate of $2.10 per reaction is a market benchmark for use with Ginkgo DSB lysate and is excluded from the 2xYTPG totals [42].

These products do not reproduce identical folding environments. GeneFrontier DnaK Mix includes GrpE in addition to DnaK/DnaJ, and DsbC Set supplies DsbC and oxidized glutathione without Erv1p; its PDI Set uses Ero1α rather than Erv1p [39]. Individual ProSpec DsbC likewise omits the remaining oxidative-folding machinery [41]. Equal folding-factor activity, optimal dose, and target-protein yield across these alternatives have not been established. Catalog leads for Erv1p, ClpB, IbpA, and IbpB lack sufficiently qualified price, active-dose, or usable-package information in the model, so they are not fully costed. Their omission does not imply that purified alternatives are unavailable or that any subset of the experimental conditions is impossible to reproduce.

The screening design comprises 6 vtPA feasibility entries, 36 vtPA DoE runs including replicated design points, and 8 Colicin M screening entries. These 50 entries do not represent 50 unique conditions or the full number of experimental reactions. For a standardized illustration, 50 reactions at the lead vtPA loading would consume $9.71 in known CFAI materials versus $430.24 for 2xYTPG with both purchased sets at the selected large-package prices. These figures are not the total expenditure of the reported campaign. Optional FluoroTect labeling alone adds $5.78 per labeled reaction and is excluded from production comparisons.

At larger final reaction volumes, recurring purchases of purified supplements produce a proportionally larger absolute cost penalty (Table S5 and Figure S3). At the standard 33.3% extract loading (5 µL per 15 µL reaction), linear projection to 1 L gives known-material costs of $12,774 for enriched CFAI and $573,268 for 2xYTPG with both purchased sets, an additional $560,494. These projections use the 50 mL-capacity supplement package prices and assume full package utilization without further bulk discounts. The relative cost difference remains constant under linear scaling. Formulation, usable extract yield, unit prices, and functional performance are assumed unchanged; preparative-scale productivity was not tested.

## 3.0 Conclusions and Outlook

Cell-free protein synthesis offers an open, directly addressable reaction environment, but its practical reach has been constrained on two fronts: extract preparation has been labor-intensive, and conventional *E. coli* extracts neither support efficient disulfide bond formation nor prevent the aggregation of folding-sensitive targets. The CFAI workflow development resolved the first constraint, and this work closes the second gap by treating the extract itself as the modular reagent so that translational capacity, chaperone identity, and oxidative folding dose can be tuned independently of the chassis strain in which each component was produced.

Two traditionally difficult and therapeutically relevant targets establish the viability and utility of the strategy. Colicin M, which lacks disulfide bonds and therefore isolates chaperone capacity from redox chemistry, was recovered as >95% soluble and active protein, showing that protein aggregation in CFAI-derived *E. coli* extracts is a tunable property of extract composition rather than a fixed limitation of the chassis. vtPA, which requires nine disulfide bonds, additionally demanded an oxidative folding capability. This work is one of the first to demonstrate that Erv1p-based oxidative folding system is both compatible with- and highly effective in *E. coli-*based cell-free systems. Across the oxidative folding conditions tested, chaperone fraction rather than total extract loading governed soluble yield, and activity measurements showed that the choice of isomerase determines functional output independently of solubility. These findings suggest that solubility may therefore be a necessary but insufficient readout, and provide additional evidence for the field’s observations that tuning a productive folding environment is a nuanced effort. Extract mixing therefore provides a practical route to screening folding environments, with its modeled materials costs considered alongside soluble yield and functional activity.

CFAI lysate mixing provides a modeled reagent-cost advantage, while the yield and activity data support a production-cost advantage by demonstrating improved soluble, functional output at comparable modeled materials inputs. The lead 50:50 Erv1p–DsbC:DnaK/J/GroESL condition at 40% extract loading combines a known-material subtotal of approximately $0.194 per 15 µL reaction with the highest measured mean vtPA activity among the three validated compositions. Colicin M likewise showed improved soluble expression with retained antimicrobial activity.

Thus, the economic relevance of mixing rests on productive protein output as well as avoided supplement purchases. Absolute costs per milligram or calibrated unit of activity, and a production-cost advantage over experimentally matched purchased -supplement workflows, remain to be established. Banking and reusing productive extract compositions can make subsequent optimization accessible to laboratories already preparing their own extracts.

Continued work will establish the generalizability of this approach. Integration with automation could extend the modeled materials-cost advantage to larger DoE campaigns and active-learning approaches that iteratively propose extract compositions. Measuring usable extract yields for each enriched strain, calibrating target-protein yield and activity, and accounting for labor and downstream processing will allow the production-cost benefit to be quantified more fully.

Demonstrating sustained functional output at preparative reaction volumes will be necessary to establish modular extract mixing as a biomanufacturing route.

## 4.0 Methods

### 4.1 CFAI Media Preparation and Culture Growth

CFAI media were prepared according to the previously described Cell-Free Autoinduction (CFAI) workflow [3, 4, 5]. The published CFAI media formulation and culture-growth procedure were followed without modification. No antibiotics were added to the CFAI growth medium.

*Escherichia coli* strains were inoculated directly from freshly streaked agar plates into CFAI medium without preparation of an overnight seed culture. Cultures were grown at 30 °C with continuous shaking at 200 rpm and harvested the following morning upon reaching stationary phase, consistent with the previously described CFAI workflow [3, 4, 5].

### 4.2 Cell Harvest and Washing

Cells were harvested and washed according to the previously described CFAI extract-preparation workflow [3, 4, 5]. All buffers and cell suspensions were maintained on ice throughout processing. Cultures were centrifuged at 5,000 × *g* at 10 °C for 10 min, and the resulting cell pellets were resuspended in 30–40 mL of pre-chilled S30 buffer prepared as previously described [5]. Following a second wash and centrifugation under identical conditions, the wet cell pellets were weighed, flash-frozen in liquid nitrogen, and stored at −80 °C until extract preparation.

Two sets of extracts were prepared to accommodate the different folding requirements of Colicin M and vtPA. For Colicin M expression, standard BL21*DE3 and chaperone-enriched extracts (GrpE/ClpB, DnaK/J/GroESL, and IbpA/B) were prepared with DTT according to the standard CFAI extract-preparation workflow [5]. For experiments involving the disulfide-bonded protein vtPA, a separate set of standard BL21*DE3 and chaperone-enriched extracts was prepared in which DTT was omitted from the S30 buffer and subsequent extract preparation. Erv1p–DsbC- and Erv1p–PDI-enriched extracts were likewise prepared without DTT. Thus, all extracts used for vtPA expression were prepared under DTT-free conditions.

### 4.3 Preparation of Cell-Free Extracts

Frozen cell pellets were thawed on ice and resuspended in S30 buffer at a ratio of 1 mL of buffer per 1 g of wet cell mass according to the previously described CFAI extract-preparation workflow [3, 4, 5]. Cell suspensions were vortexed until homogeneous and lysed by sonication at 20 kHz and 50% amplitude in 1.4 mL aliquots using three 45 s sonication cycles separated by 59 s cooling intervals on ice [5]. Following lysis, crude extracts were clarified by centrifugation at 18,000 × *g* at 4 °C for 10 min. The resulting supernatants were collected as functional cell-free extracts, aliquoted, flash-frozen in liquid nitrogen, and stored at −80 °C until use. Extracts used for Colicin M and vtPA expression were prepared as described above, following either the standard DTT-containing or DTT-free CFAI workflow, respectively.

### 4.4 Plasmid Construct and Template Preparation

Chaperone-encoding plasmids pBB540 (GrpE/ClpB; Addgene plasmid #27393; RRID: Addgene_27393), pBB550 (DnaK/DnaJ/GroESL; Addgene plasmid #27396; RRID: Addgene_27396), and pBB572 (IbpA/B; Addgene plasmid #27397; RRID: Addgene_27397) were gifts from Bernd Bukau and obtained from Addgene. Oxidative-folding plasmids pCOLA-Gent-EM7-Erv1p–DsbC (Addgene plasmid #202485; RRID: Addgene_202485) and pCOLA-Gent-EM7-Erv1p–PDI (Addgene plasmid #202486; RRID: Addgene_202486) were gifts from Michael Lynch and obtained from Addgene. The oxidative-folding plasmids constitutively encode the sulfhydryl oxidase Erv1p together with either the disulfide isomerase DsbC or protein disulfide isomerase (PDI), respectively.

Target-protein expression constructs encoding Colicin M and a truncated variant of human tissue plasminogen activator (vtPA) were provided by the Jewett laboratory (Stanford University). Both target constructs were supplied in the pJL1 backbone for T7-driven cell-free expression and were used as received. Exact construct boundaries were not independently determined or sequence-verified in this study.

Chaperone and oxidative-folding plasmids were transformed into *Escherichia coli* BL21*(DE3), whereas target-protein expression plasmids were propagated in NEB 5-alpha. Bacterial transformations were performed by heat shock according to the Addgene bacterial transformation protocol [43]. Following heat shock, cells were recovered in SOC medium at 37 °C with shaking for 45 min and plated on LB agar containing the appropriate antibiotic corresponding to the resistance marker of each plasmid. Plasmid DNA was purified using a Zymo Research plasmid miniprep kit, eluted in nuclease-free water, and quantified using a NanoDrop spectrophotometer.

### 4.5 Cell-Free Protein Synthesis (CFPS) Reaction Assembly and Optimization

Standard 15 µL reactions were assembled in sterile 1.5 mL microcentrifuge tubes at a fixed total extract concentration of 33% v/v (5.0 µL), unless otherwise noted, and incubated at 30 °C for 20 hours. Each reaction contained 16 ng/µL plasmid template and a reagent mixture prepared exactly as described previously [5], with no modifications to the published PANOx-SP formulation.

### 4.6 Combinatorial Mixing Strategy

To tune the folding environment, the extract component of each reaction was composed of modular mixtures of specialized extracts. Mixtures were assembled volumetrically immediately prior to reaction assembly and were not pre-incubated.

#### Feasibility screen

Initial screening used a fixed 50:50 volumetric ratio of one oxidative folding extract (Erv1p-PDI or Erv1p-DsbC) to one chaperone extract (pBB540, pBB550, or pBB572), at a total extract concentration of 33% v/v. Solubility was assessed by SDS-PAGE as described below.

#### Design of Experiments

For vtPA, a full factorial design investigated two continuous factors: the oxidative folding-to-chaperone volumetric ratio (0:100, 25:75, 50:50, 75:25, and 100:0) and the total extract concentration (25%, 33%, and 40% v/v), yielding 15 unique design points. Three additional replicate runs were included to estimate pure error: two replicates of the center point (50:50 ratio at 33% v/v, Condition 8) and one replicate of the 0:100 condition at 33% v/v. Each replicate was prepared as a wholly independent reaction. Runs were performed in the order listed in Table 2. The identical design was applied independently to the Erv1p-DsbC and Erv1p-PDI systems, each paired with the DnaK/J/GroESL chaperone extract.

### 4.7 SDS-PAGE and Densiometric Quantification

Total and soluble fractions were diluted 1:10 by combining 2 µL of each fraction with 18 µL of ultrapure water. For reducing conditions, 9 µL of diluted sample was combined with 1 µL of 1 M DTT and 2 µL of 6× Laemmli buffer and denatured at 95 °C for 5 minutes. For non-reducing conditions, DTT was omitted and samples were denatured under identical conditions. Samples were separated on hand-cast 12% polyacrylamide gels. Resolving gels (12%) were prepared with 8.8 mL water, 5 mL 1.5 M Tris-HCl pH 8.8, 200 µL 10% SDS, 6 mL 40% acrylamide, 200 µL 10% APS, and 10 µL TEMED. Stacking gels (4%) were prepared with 6.4 mL water, 2.5 mL 0.5 M Tris-HCl pH 6.8, 100 µL 10% SDS, 1 mL 40% acrylamide, 100 µL 10% APS, and 5 µL TEMED. Electrophoresis was performed in 1× Tris-glycine-SDS running buffer at 180 V for 30 to 40 minutes.

Colicin M gels were stained with Coomassie Brilliant Blue R-250 for 45–60 min and subsequently destained for 120–180 min using a destaining solution consisting of 50% (v/v) methanol, 10% (v/v) glacial acetic acid, and 40% (v/v) deionized water. Gels were imaged using a Bio-Rad Gel Doc imaging system (Bio-Rad Laboratories). Densitometric analysis was performed using ImageJ. For each lane, a rectangular region of interest (ROI) was drawn around the band corresponding to the target protein, and integrated density was measured. Local background was determined using an equivalently sized ROI positioned adjacent to the target band within the same lane. Background-corrected integrated density was calculated by subtracting the integrated density of the background ROI from that of the corresponding target-band ROI.

Integrated density values are reported in arbitrary units. For the Colicin M experiments, values were normalized to the non-supplemented BL21*(DE3) control. For the Design of Experiments analysis, values were normalized within each oxidative folding system to its respective center-point replicate (Condition 8), assigned a reference value of 100 A.U. Percent solubility was calculated as the ratio of soluble to total integrated density for each condition.

### 4.8 Fluor0Tect Labeling of vtPA

vtPA was labeled co-translationally by including 0.33 µL of FluoroTect GreenLys tRNA (Promega, L5001) per 15 µL cell-free reaction. Following incubation and separation of total and soluble fractions, samples were prepared as described above and resolved by SDS-PAGE. FluoroTect-labeled vtPA was visualized directly in the gel using the GelGreen fluorescence imaging setting on a Bio-Rad Gel Doc imaging system.

### 4.9 sfGFP Quantification

sfGFP expression was quantified by fluorescence. Following incubation, 2 µL of each cell-free reaction was diluted with 48 µL of 0.5 M HEPES in a black 96-well half-area plate.

Fluorescence was measured using a plate reader with excitation and emission wavelengths of 485 and 510 nm. Fluorescence values are reported as relative fluorescence units (RFU) without conversion to absolute protein concentration.

### 4.10 Growth Curve Measurement

Culture density was monitored at 30–60 min intervals by removing 1 mL aliquots, diluting samples as necessary to remain within the linear range of the spectrophotometer, and measuring absorbance at 600 nm (OD600). Samples exceeding an OD600 of 1 were diluted 10-fold in Luria Broth prior to measurement and corrected accordingly. Growth curves represent n = 7 biological replicates.

### 4.11 Colicin M Activity Assay

Antimicrobial activity of cell-free-expressed Colicin M was evaluated using an *Escherichia coli* K-12 cell-killing assay. *E. coli* K-12 cultures were grown to mid-log phase (OD600 ≈ 0.8), harvested by centrifugation at 3,000 × *g* for 3 min, and resuspended in Luria broth (LB) to an OD600 of 0.1. For each treatment, 50 µL of the resulting cell suspension was combined with 2.5, 7.5, or 12.5 µL of Colicin M-containing CFPS reaction. The total volume of CFPS material was maintained at 12.5 µL by supplementation with no-DNA CFPS control reaction, corresponding to 10, 5, and 0 µL of no-DNA CFPS for the 2.5, 7.5, and 12.5 µL Colicin M treatments, respectively. Cells-only and 12.5 µL no-DNA CFPS controls were included.

Samples were incubated for 1 h at 37 °C with shaking at 220 rpm. Following incubation, samples were plated directly onto LB agar and incubated overnight at 37 °C. Viable colonies were counted and reported as colony-forming units (CFU).

### 4.12 vtPA Activity Assay

vtPA protease activity was assessed using a fluorogenic substrate assay. Cell-free reactions were incubated at 30 °C for approximately 20 h and subsequently centrifuged at 5,000 × *g* for 10 min at 4 °C to remove insoluble material. The resulting soluble fractions were collected and diluted 5-fold prior to activity measurements.

For each assay, 10 µL of diluted soluble fraction was combined with 40 µL of substrate solution containing 50 µM D-Ile-Pro-Arg-AMC (Echelon Biosciences, 855-18) in 100 mM Tris-HCl (pH 8.8) containing 0.01% (v/v) Tween-20 in a black 96-well half-area microplate. Release of fluorescent 7-amino-4-methylcoumarin (AMC) was monitored using a Cytation microplate reader at excitation and emission wavelengths of 354 and 442 nm, respectively. Fluorescence was measured over 1 h at 37 °C with intermittent shaking.

Protease activity was quantified as the slope of the linear portion of each fluorescence-versus-time trace. Slopes were corrected by subtracting contributions from both a no-DNA CFPS control and a substrate-only background. Because a purified vtPA standard was not included, activity is reported as relative fluorescence units per minute (RFU/min) rather than as absolute enzymatic activity.

### 4.13 Statistical Analysis and Response-Surface Modeling

Unless otherwise noted, cell-free reactions were performed in n = 3 triplicates, defined as independent reactions. Error bars in Figures 1, 2, and 3 represent the standard deviation across replicates. Error bars in Figure 7 represent the standard error of the mean for n = 3 replicates. The specific replicate structure for each experiment is stated in the corresponding figure caption.

For the Design of Experiments, each of the 15 unique design points was performed once. Three additional runs were included to estimate pure error: two further runs at the center point (50:50 ratio at 33% v/v, Condition 8) and one further run at the 0:100 condition at 33% v/v, giving 18 runs in total. Each run was prepared as a wholly independent reaction rather than a repeated measurement of a single reaction.

Normalized soluble vtPA integrated density was used as the response variable for response-surface modeling rather than percent solubility, as the latter is bounded at >95% and saturates in the high-performing region, obscuring differences in absolute yield. A second -order response-surface model was fit separately for each oxidative folding system, incorporating main effects for total extract loading and chaperone ratio, their two-factor interaction, and quadratic terms for both factors. Both factors were centered and scaled prior to fitting, so coefficients are reported in coded units. Models were fit by ordinary least squares in MATLAB (MathWorks) using base functionality only. Coefficient standard errors and t-statistics were computed from the ordinary least-squares covariance matrix, and two-sided p-values and the overall model F-test from the regularized incomplete beta function. Overall model significance was assessed by F-test against a constant model, and individual coefficient significance by t-test, with p < 0.05 considered significant. Reported R² and adjusted R² values refer to the full second -order model. Pure error was estimated from the replicated factor settings and lack of fit was tested against it. Predicted R² was computed from the leave-one-out predicted residual error sum of squares (PRESS), and adequate precision was calculated as the range of predicted responses divied by the mean predicttion standard error. Response surfaces were rendered as filled contour plots in MATLAB, with predicted values evaluated from the fitted coefficients across a grid spanning the experimental design space. Contours represent model predictions rather than interpolation between observations, and the 18 experimental design points are overlaid as markers.

Specific activity values for vtPA were calculated as the slope of the linear portion of each kinetic trace, corrected by subtraction of a no-DNA CFPS control and a substrate-only background.

Uncertainty in the corrected slope was determined by propagating the standard errors of the sample, no-DNA control, and substrate-only background slopes in quadrature, and is reported as propagated standard error of the mean.

### 4.14 Safety

Colicin M is a bacteriocin with antimicrobial activity against susceptible *E. coli* strains. Cell-free reactions containing Colicin M and cultures used in cell-killing assays were handled under BSL-1 containment and decontaminated with bleach prior to disposal. No other unexpected or significant hazards were associated with this work.

## AUTHOR INFORMATION

### Author Contributions

L.F. and A.D. designed and performed the experiments, analyzed the data, and wrote the manuscript. J.P.O. conceived and supervised the project, secured funding, and edited the manuscript. All authors have given approval to the final version of the manuscript.

## ACKNOWLEDGMENTS

We thank the Dr. Megan Olsen and the Jewett laboratory (Stanford University) for providing the Colicin M and vtPA expression constructs. We thank the William and Linda Frost Center for Research and Innovation at California Polytechnic State University, San Luis Obispo, for providing access to the research facilities in which this work was conducted.

## Supplemental Data

**Figure S1.**
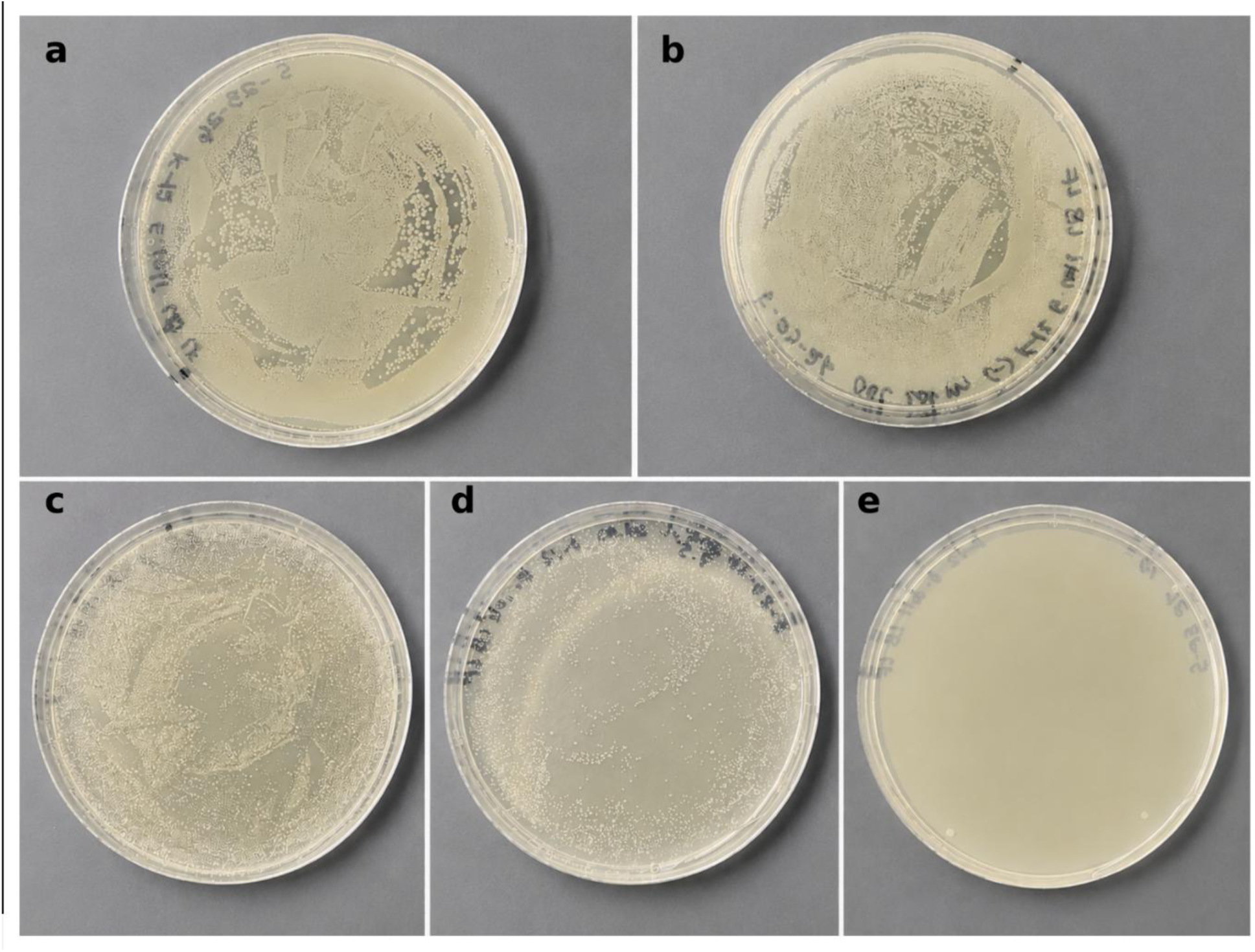
Functional validation of cell-free expressed Colicin M by cell-killing assay. Mid-log phase *E. coli* K-12 cells were incubated with increasing volumes of CFPS-derived Colicin M extract (2.5, 7.5, or 12.5 µL). To maintain a constant CFPS volume, negative-control CFPS extract lacking the Colicin M expression plasmid was added as needed. Representative plates from one of three independent biological replicates are shown. Plate **(a)**: K-12 cells only (0 µL CFPS). Plate **(b)**: K-12 cells treated with 12.5 µL negative-control CFPS extract. Plates **(c–e)**: K-12 cells treated with 2.5, 7.5, and 12.5 µL Colicin M CFPS extract, respectively. Control cells and cells treated with negative-control CFPS exhibited confluent growth, while increasing volumes of Colicin M extract resulted in progressively reduced colony formation, culminating in near-complete elimination of viable colonies at 12.5 µL. Colony counts were determined from three independent biological replicates.

