## Supplemental Materials for "Modular Cell-Free Extract Mixing Enables Soluble, Active Biomanufacturing of Difficult-to-Express and Aggregation-Prone Proteins"

**Figure S1. Functional validation of cell-free expressed Colicin M by cell-killing assay.**

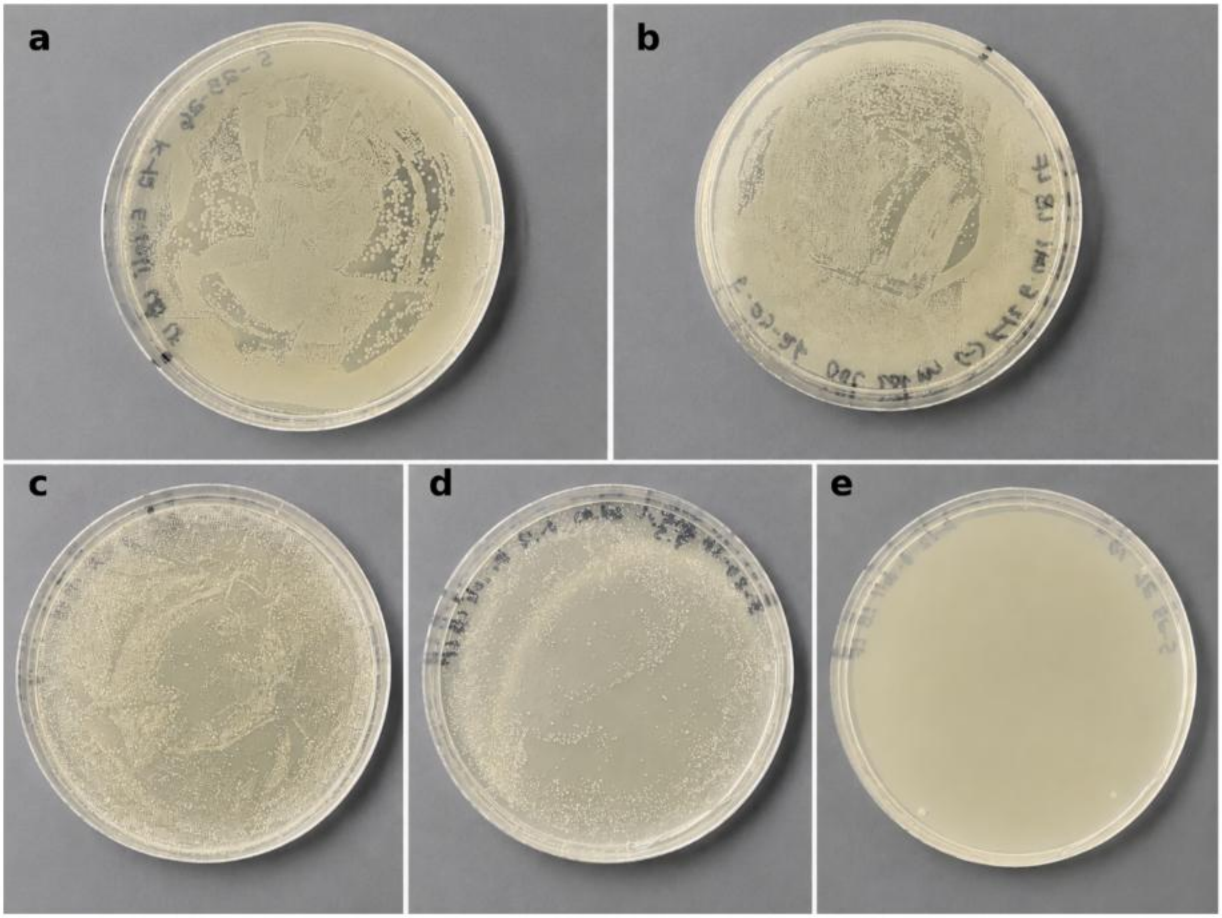

**Figure S1.** Functional validation of cell-free expressed Colicin M by cell-killing assay. Mid-log phase E. coli K-12 cells were incubated with increasing volumes of CFPS-derived Colicin M extract (2.5, 7.5, or 12.5 µL). To maintain a constant CFPS volume, negative-control CFPS extract lacking the Colicin M expression plasmid was added as needed. Representative plates from one of three independent biological replicates are shown. Plate (a): K-12 cells only (0 µL CFPS). Plate (b): K-12 cells treated with 12.5 µL negative-control CFPS extract. Plates (c–e): K-12 cells treated with 2.5, 7.5, and 12.5 µL Colicin M CFPS extract, respectively. Control cells and cells treated with negative-control CFPS exhibited confluent growth, while increasing volumes of Colicin M extract resulted in progressively reduced colony formation, culminating in near-complete elimination of viable colonies at 12.5 µL. Colony counts were determined from three independent biological replicates.

Table S1. Full factorial design and observed responses. Total extract loading and chaperone fraction are the two factors; L and C are the corresponding coded values, L = (loading – 33)/15 and C = (chaperone fraction – 50)/100. Responses are soluble vtPA from densitometry, normalized to the reference condition.

| Run | Total Extract Loading (%v/v) | Chaperone Fraction (%) | L (coded) | C (coded) | Erv1p-DsbC (norm A.U.) | Erv1p-PDI (norm A.U.) | |
|---|---|---|---|---|---|---|---|
| 1 | 25 | 100 | -0.5333 | 0.50 | 0.8911 | 1.0844 | |
| 2 | 25 | 75 | -0.5333 | 0.25 | 1.0268 | 1.3465 | |
| 3 | 25 | 50 | -0.5333 | 0.00 | 0.4614 | 0.6149 | |
| 4 | 25 | 25 | -0.5333 | -0.25 | 0.1453 | 0.5442 | |
| 5 | 25 | 0 | -0.5333 | -0.50 | 0.0023 | 0.0612 | |
| 6 | 33 | 100 | 0.0000 | 0.50 | 0.6057 | 1.7095 | a |
| 7 | 33 | 75 | 0.0000 | 0.25 | 0.6453 | 1.1876 | |
| 8 | 33 | 50 | 0.0000 | 0.00 | 0.7671 | 0.8606 | b |
| 9 | 33 | 25 | 0.0000 | -0.25 | 0.4283 | 0.2151 | |
| 10 | 33 | 0 | 0.0000 | -0.50 | 0.2669 | 0.1669 | |
| 11 | 40 | 100 | 0.4667 | 0.50 | 0.6657 | 0.8857 | |
| 12 | 40 | 75 | 0.4667 | 0.25 | 1.2900 | 1.1857 | |
| 13 | 40 | 50 | 0.4667 | 0.00 | 1.0995 | 0.9831 | |
| 14 | 40 | 25 | 0.4667 | -0.25 | 0.6908 | 1.3284 | |
| 15 | 40 | 0 | 0.4667 | -0.50 | 0.5646 | 0.2303 | |
| 16 | 33 | 50 | 0.0000 | 0.00 | 0.6029 | 0.6567 | b |
| 17 | 33 | 50 | 0.0000 | 0.00 | 0.6652 | 0.8081 | b |
| 18 | 33 | 100 | 0.0000 | 0.50 | 0.6989 | 1.0121 | a |

a *Replicate of the 33% loading / 100% chaperone setting (n = 2).* b *Replicate of the 33% loading / 50% chaperone setting (n = 3). These replicated settings provide the 3 degrees of freedom of pure error used for the lack-of-fit test in Table S2.*

Table S2. Analysis of variance for the quadratic response surface models of soluble vtPA. Term sums of squares are partial (Type III). Lack of fit is tested against pure error estimated from the replicated factor settings identified in Table S1.

| | Erv1p-DsbC | | | | | Erv1p-PDI | | | | |
|---|---|---|---|---|---|---|---|---|---|---|
| **Source** | **SS** | **df** | **MS** | **F** | **p** | **SS** | **df** | **MS** | **F** | **p** |
| Model | 1.3955 | 5 | 0.2791 | 9.577 | 0.0007 | 2.5012 | 5 | 0.5002 | 5.487 | 0.0074 |
| L | 0.3782 | 1 | 0.3782 | 12.978 | 0.0036 | 0.1088 | 1 | 0.1088 | 1.193 | 0.2961 |
| C | 0.6643 | 1 | 0.6643 | 22.794 | 0.0005 | 2.1657 | 1 | 2.1657 | 23.754 | 0.0004 |
| L × C | 0.1791 | 1 | 0.1791 | 6.146 | 0.0290 | 0.1282 | 1 | 0.1282 | 1.406 | 0.2587 |
| $L^2$ | 0.1152 | 1 | 0.1152 | 3.954 | 0.0701 | 0.0302 | 1 | 0.0302 | 0.331 | 0.5756 |
| $C^2$ | 0.2259 | 1 | 0.2259 | 7.750 | 0.0165 | 0.1504 | 1 | 0.1504 | 1.649 | 0.2233 |
| Residual | 0.3497 | 12 | 0.0291 | — | — | 1.0940 | 12 | 0.0912 | — | — |
| Lack of fit | 0.3316 | 9 | 0.0368 | 6.114 | 0.0818 | 0.8284 | 9 | 0.0920 | 1.039 | 0.5483 |
| Pure error | 0.0181 | 3 | 0.0060 | — | — | 0.2657 | 3 | 0.0886 | — | — |
| Cor total | 1.7452 | 17 | — | — | — | 3.5953 | 17 | — | — | — |
| **$R^2$** | 0.7996 | | | | | 0.6957 | | | | |
| **Adjusted $R^2$** | 0.7161 | | | | | 0.5689 | | | | |
| **Predicted $R^2$** | 0.4895 | | | | | 0.2026 | | | | |
| **PRESS** | 0.8909 | | | | | 2.8667 | | | | |
| **Std. deviation** | 0.1707 | | | | | 0.3019 | | | | |
| **Mean** | 0.6399 | | | | | 0.8267 | | | | |
| **C.V. (%)** | 26.68 | | | | | 36.52 | | | | |
| **Adequate precision** | 11.20 | | | | | 7.83 | | | | |

*For both systems, the model was significant, and lack of fit was not significant at the 0.05 level, and adequate precision exceeded 4, indicating sufficient signal to resolve the response across the design space. Pure error is estimated from only 3 degrees of freedom (two replicated settings), so the lack-of-fit test has limited power to detect model inadequacy and the result should be read as an absence of evidence against the model rather than as strong evidence for it. For both systems the predicted $R^2$ differs from the adjusted $R^2$ by more than 0.20 (0.227 for Erv1p–DsbC, 0.366 for Erv1p–PDI), indicating limited predictive capability beyond the measured design points; the fitted surfaces are therefore interpreted as descriptive of trends within the tested range rather than as predictive optimization models.*

Table S3. Regression coefficient estimates for the quadratic response surface models, in coded factor units.

| | Erv1p–DsbC | | | | Erv1p–PDI | | | |
|---|---|---|---|---|---|---|---|---|
| **Term** | **Coefficient** | **Std. error** | **t** | **p** | **Coefficient** | **Std. error** | **t** | **p** |
| Intercept | 0.6639 | 0.0729 | 9.103 | <0.0001 | 0.8564 | 0.1290 | 6.638 | <0.0001 |
| L | 0.4002 | 0.1111 | 3.603 | 0.0036 | 0.2147 | 0.1965 | 1.092 | 0.2961 |
| C | 0.5672 | 0.1188 | 4.774 | 0.0005 | 1.0242 | 0.2101 | 4.874 | 0.0004 |
| L × C | -0.7564 | 0.3051 | -2.479 | 0.0290 | -0.6399 | 0.5397 | -1.186 | 0.2587 |
| $L^2$ | 0.6524 | 0.3281 | 1.989 | 0.0701 | 0.3340 | 0.5803 | 0.575 | 0.5756 |
| $C^2$ | -1.0455 | 0.3756 | -2.784 | 0.0165 | -0.8531 | 0.6643 | -1.284 | 0.2233 |

*Coefficients are reported in coded units so that their magnitudes are directly comparable. Fitted model, Erv1p–DsbC:* $\hat{y} = 0.6639 + 0.4002\,L + 0.5672\,C - 0.7564\,LC + 0.6524\,L^2 - 1.0455\,C^2$. *Erv1p–PDI:* $\hat{y} = 0.8564 + 0.2147\,L + 1.0242\,C - 0.6399\,LC + 0.3340\,L^2 - 0.8531\,C^2$.

### Supplementary Cost Analysis

We estimated known materials consumed by six cell-free protein synthesis routes using 15 µL reactions with 5 µL (33.3% v/v) in-house-produced extract. Each route includes priced PANOx reagents and an extract-preparation allocation. CFAI with both enriched folding systems also includes allocated strain and construct acquisition. The purchased-supplement cases use 2xYTPG extract with GeneFrontier DnaK/GroE chaperones, a DsbC-based disulfide set, or both, at the dose and 50 mL-capacity pack tier recorded in the cost workbook.

The model assumes 10 mL usable CFAI extract per liter of culture and equal recovery from enriched strains. The 2xYTPG comparison uses a 4.29 mL/L literature proxy. At the selected pack tier, purchased chaperones contribute $6.5556 and the disulfide set contributes $1.8354 per 15 µL reaction. Vendor products have different compositions and compatibility requirements; the graph does not treat them as interchangeable. Dose and active-protein yields were not matched experimentally.

**Table S4.** Known-material cost per 15 µL reaction with 5 µL in-house extract. The supplement column combines the purchased chaperone and disulfide components when applicable. Values are consumption estimates in USD.

| Route | Reagents + extract | Purchased supplements | Total |
|---|---:|---:|---:|
| CFAI, not enriched | $0.1872 | $0.0000 | $0.1872 |
| CFAI, both enriched systems | $0.1916 | $0.0000 | $0.1916 |
| 2xYTPG, no supplements | $0.2083 | $0.0000 | $0.2083 |
| 2xYTPG + disulfide set | $0.2080 | $1.8354 | $2.0434 |
| 2xYTPG + chaperones | $0.2083 | $6.5556 | $6.7639 |
| 2xYTPG + both sets | $0.2080 | $8.3910 | $8.5990 |

**Table S5.** Linear projection of known-material costs at larger final reaction volumes. Large-pack supplement unit costs are held constant and fully used; no bulk discount or scale-dependent change in extract recovery is assumed. Values are rounded to the nearest dollar.

| Final volume | Enriched CFAI | 2xYTPG + both sets | Additional cost |
|---|---:|---:|---:|
| 50 mL | $639 | $28,663 | $28,025 |
| 100 mL | $1,277 | $57,327 | $56,049 |
| 500 mL | $6,387 | $286,634 | $280,247 |
| 1 L | $12,774 | $573,268 | $560,494 |

At 1 L, the calculation represents 66,666.7 reaction equivalents and uses twenty 50 mL pack equivalents for each purchased supplement product. GeneFrontier product listings supplied the price basis recorded in the workbook (https://purefrex.genefrontier.com/products/). These comparisons omit unpriced template DNA and preparation materials, as well as labor, equipment, infrastructure, and losses not specified in the model.

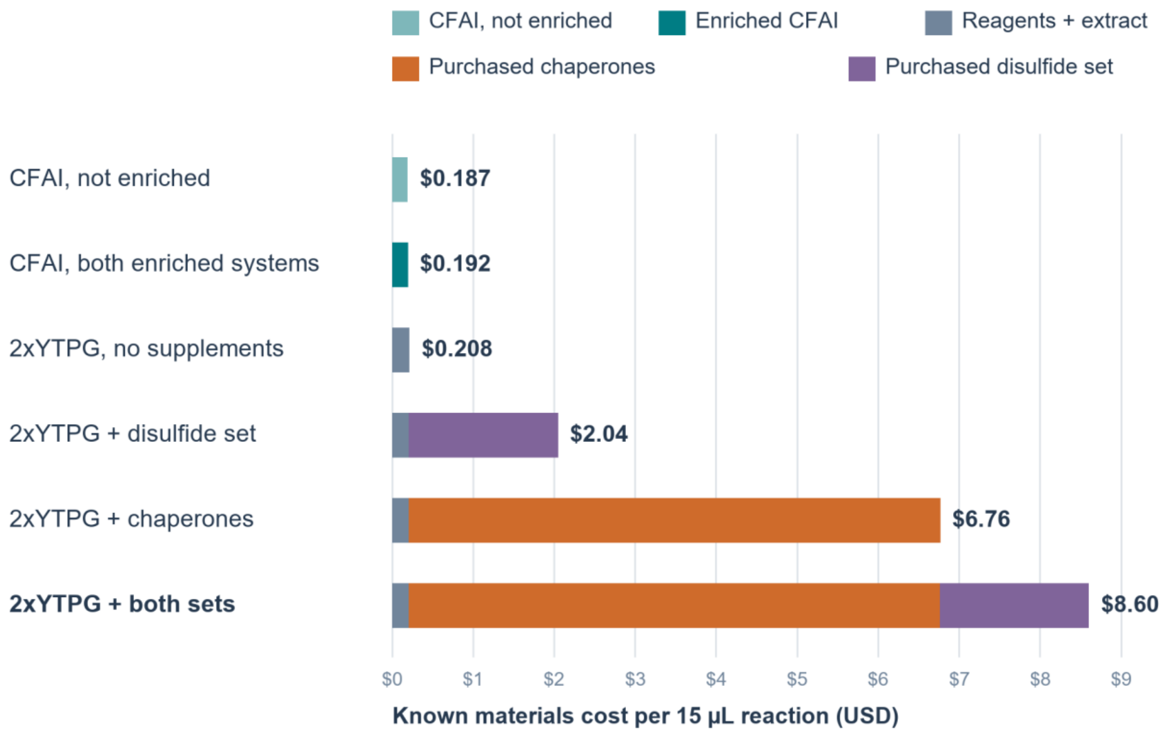


**Figure S2.** Known-material cost per 15 µL reaction at 5 µL (33.3% v/v) in-house extract loading. CFAI without enrichment costs $0.1872 and CFAI with both enriched folding systems costs $0.1916 per reaction. The 2xYTPG route costs $0.2083 without purchased folding supplements, $2.0434 with the disulfide set, $6.7639 with chaperones, and $8.5990 with both sets. Purchased supplements account for $8.3910 of the both-set total at the GeneFrontier large-pack unit price. These are cost scenarios, not activity-matched substitutes or estimates of cost per active protein.

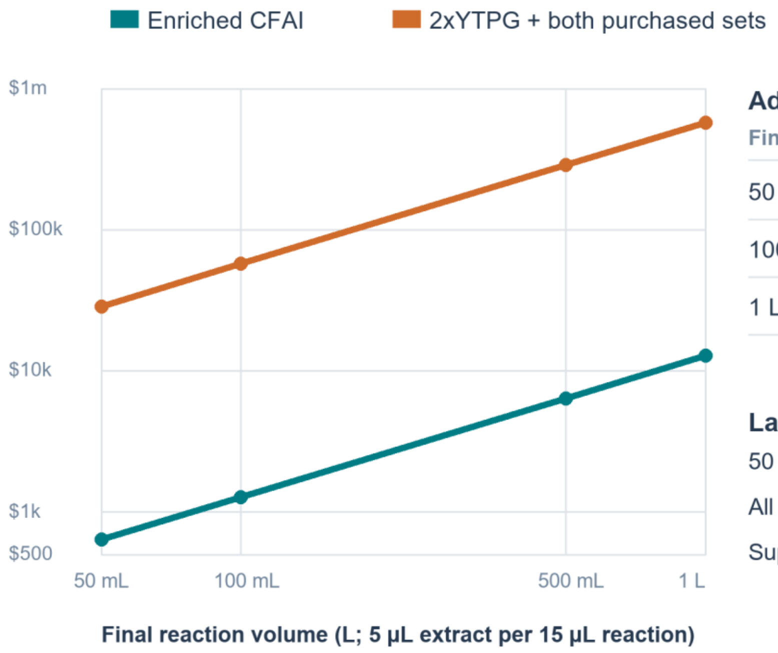


#### Additional cost vs enriched CFAI

| Final volume | Large-pack case |
|---|---|
| 50 mL | **$28,025** |
| 100 mL | **$56,049** |
| 1 L | **$560,494** |

#### Large-pack supplement basis at 1 L

| | |
|---|---|
| 50 mL pack equivalents | 20 |
| All three sets per 50 mL | $27,970 |
| Supplement cost per 1 L | **$559,400** |

**Figure S3.** Projected known-material costs for enriched CFAI and 2xYTPG with both purchased sets from 50 mL to 1 L of final reaction volume. The projection multiplies the per-15 µL costs in Table S4 by the number of reaction equivalents. At 1 L, enriched CFAI costs $12,774 and the supplemented 2xYTPG route costs $573,268, an additional $560,494. The supplement component alone is $559,400 at the 50 mL pack tier, repeated twenty times with full use and no bulk discount. Formulation, recovery, unit prices, and functional performance at scale were not validated by this calculation.
